# Chromosomal instability shapes spatial and temporal phenotypic diversity in a malignant peripheral nerve sheath tumour

**DOI:** 10.64898/2026.08.13.744699

**Authors:** Haixi Yan, Jonas Demeulemeester, Annelien Verfaillie, Yixiao Cheng, Yidan Pan, Cristina Cotobal Martin, Sophia Ward, Alexander Stein, Tom Lesluyes, Tom Kaufmann, Roland F. Schwarz, Thierry Voet, Charles Swanton, Simone Zaccaria, Adrienne M. Flanagan, Maxime Tarabichi, Peter Van Loo

## Abstract

Understanding the spatial and temporal dynamics of tumour evolution is crucial for determining the drivers of cancer progression. Linking genotype to phenotype across a tumour remains challenging, however. Here, we use single-cell and spatial multi-omics to comprehensively profile a primary malignant peripheral nerve sheath tumour (MPNST) and its multifocal recurrence. Combining the native barcoding system from extensive heterogeneity in copy number alterations with mutation data, we resolve the evolutionary tree of this tumour, revealing a branching structure suggestive of ongoing chromosomal instability. We show that gene dosage effects contribute to phenotypic diversity and scale predominantly linearly with copy number. Using spatial genomics assisted by laser capture microdissection and spatial transcriptomics, we performed *in situ* lineage tracing in this human tumour, elucidating the relationship between local expansions and interactions between tumour cells and the microenvironment. These findings demonstrate the potential of combined bulk, single-cell and spatial techniques to dissect cancer evolution in time and space and link genotype to phenotype in detail.

## Introduction

Tumours are typically composed of a diverse patchwork of subclones, driving cancer evolution and disease progression^1^. This genetic heterogeneity contributes to phenotypic heterogeneity which, in turn, underpins metastasis, therapy resistance and poor prognosis^2,3^. Recent studies have begun to characterise the degree of heterogeneity within individual tumours with increasing resolution. Multi-region and representative sequencing allow for enhanced sampling of a tumour and sensitive detection of subclones, enabling the reconstruction of detailed phylogenetic trees^4^. Furthermore, single-cell approaches allow for tumour heterogeneity to be explored more comprehensively still, by characterising individual cancer cells as well as those of the microenvironment^5–7^. Spatial information has recently also been incorporated to elucidate interactions between these cells further and detail the relationship between tumour subclones^8,9^. A multi-omics approach allows for the integration of these information layers and to bridge the gap from genotype to phenotype.

Malignant peripheral nerve sheath tumours (MPNSTs) are rare, highly aggressive soft tissue sarcomas thought to be derived from Schwann cells or neural crest progenitors^10^. Treatment options are limited with surgical resection as the mainstay of treatment. However, recurrence rates are high, and prognosis is poor with a 5-year overall survival rate of 26-39%^11^. Around half of all MPNSTs occur in patients with neurofibromatosis type 1 (NF1)^12^. These patients show variable phenotypic severity and have a lifetime risk of 9-13% of developing MPNST^13^. Previous studies of MPNST have identified mutations in *NF1*, *TP53*, *CDKN2A* and the polycomb repressive complex 2 (PRC2) components *EED* and *SUZ12* as critical drivers in its pathogenesis^14^. More detailed genomic profiling of MPNSTs, although limited in scale, has shown that these tumours are genomically complex^15^. They typically exhibit a moderate burden of single nucleotide variants (SNVs) but display frequent copy number alterations (CNAs) and high ploidy.

Here, we performed deep multi-omics integration and spatial analysis of a recurrent MPNST with single-cell resolution. By leveraging the extensive heterogeneity in CNAs as a native barcoding system, we link genetic subclones to clusters of cells and their transcriptomes. We validate this approach with SNVs and genome and transcriptome (G&T) sequencing of the same cells. We explore gene dosage effects and show that these explain nearly all phenotypic diversity observed. Finally, we map genomic and phenotypic heterogeneity in 2D and 3D, showcasing extensive micro- and macroscopic diversity, growth trajectories, adaptation, and parallel evolution. Our work demonstrates the power of combining spatial multi-omics at the single-cell and bulk levels to study cancer evolution.

## Results

### Multi-region bulk and single-cell whole-genome sequencing reveals an early common ancestor of post-treatment recurrence regions

Our index case was a 36-year-old male who developed an MPNST on the right forearm. The patient exhibited an NF1 phenotype clinically and correspondingly harboured a germline *NF1* c.269T>C missense mutation (p.L90P). Despite surgical resection of the primary tumour, followed by 6 cycles of adjuvant doxorubicin and ifosfamide, the patient developed a multifocal local recurrence and underwent palliative resection through amputation of the right arm (**Fig. 1a**). We sampled the primary tumour and 5 recurrence regions and investigated intra-tumour heterogeneity using a multi-region bulk and single-cell multi-omics approach. H3K27me3 loss^16^ was detected upon histological review. This phenotype is associated with mutations in PRC2 components and correspondingly, we identified a homozygous deletion of *SUZ12*. We also identified a frameshift-causing deletion in *TP53*.

**Figure 1.**
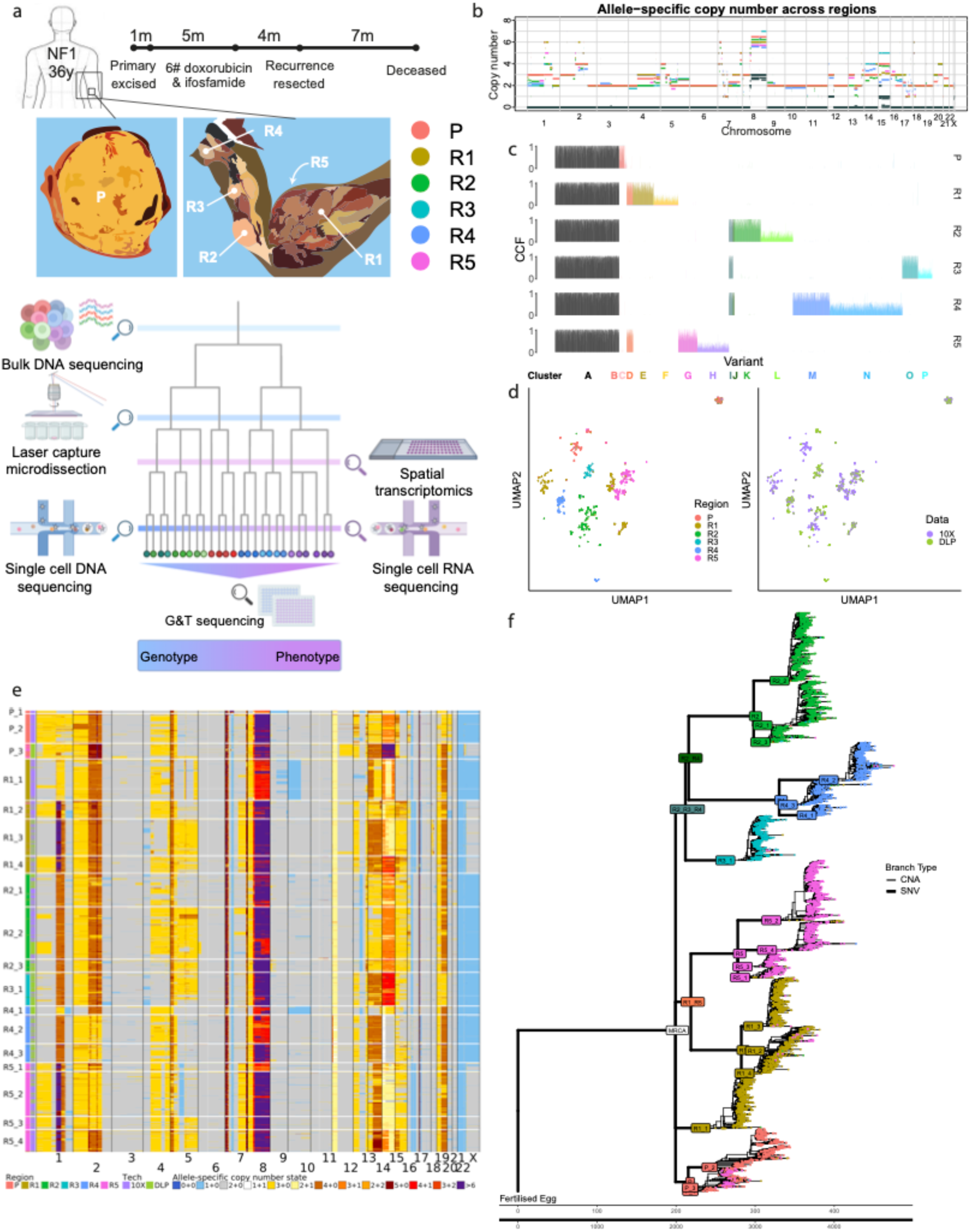
Study overview and single-cell phylogenetic reconstruction from bulk and single-cell data. (**a**) Clinical history of the patient and schematic of our multi-omic study design. The primary sample (P) was collected from the excised tumour in the forearm. Recurrence regions R1 and R5 were sampled from opposite sides of the forearm where the primary initially developed and R2, R3 and R4 were sampled from the upper arm. (**b**) Allele-specific copy number profiles from Battenberg with the total copy number coloured by sample as in panel (a) and the minor allele copy number in teal. Subclonal copy number states are shown as non-integer states proportional to size of the subclone. (**c**) CCF of SNVs in each region derived from ndDPClust with SNVs coloured by cluster. (**d**) Uniform Manifold Approximation and Projection (UMAP) of integer copy number profiles of all single-cell DNA cells annotated by region (left) and sequencing technology (right). Non-tumour cells are seen in the top right of the plots. (**e**) Allele-specific copy number profiles of tumour cells in each K-means defined subclones. (**f**) Full phylogenetic tree with number of SNVs (from ndDPClust phylogenetic reconstruction) shown by thicker branches and number of CNA events (from MEDICC2 phylogenetic reconstruction) shown with thinner lines. Most recent common ancestor (MRCA) shown in white.

We performed bulk whole-genome sequencing of all samples to a median depth of 54X (range 46-67X) and derived subclonal CNA profiles using Battenberg^17^. Fluorescence-Activated Cell Sorting (FACS) analysis revealed that this tumour had a ploidy of 2.6, while Battenberg copy number analysis highlighted near genome-wide clonal loss of heterogeneity (LOH) (>91.7%; **Fig. 1b** and **Supplementary Fig. 1**), except on chromosomes 8, 12, and 15. Therefore, this tumour likely developed into a near-haploid state before undergoing a whole-genome doubling (WGD). In addition, we consistently observed both inter-region and intra-region subclonal copy number heterogeneity (**Supplementary Fig. 2**).

Next, we called 11,734 SNVs and identified truncal, shared clonal, exclusive clonal and one subclonal mutation cluster for each region using multidimensional Bayesian Dirichlet Process-based mutation clustering (ndDPClust) (**Fig. 1c**)^18^. By assessing the cancer cell fraction (CCF) of clusters across samples and applying the pigeon-hole principle, we reconstructed an evolutionary tree for this tumour to the resolution of a single subclone for each sample (**Supplementary Table 1**, **Supplementary Fig. 3a**)^17^. This revealed that all recurrence regions were derived from a common ancestor that was present prior to primary tumour removal. While one tumour lineage seeded the recurrence regions R1 and R5, a sibling lineage gave rise to R2, R3 and R4, recapitulating their spatial proximity on the arm. We also applied MEDICC2^19^, a copy number-based maximum parsimony method, resulting in a tree confirming these early branching events of the SNV-based tree (**Supplementary Fig. 3b**). We performed fitting of SNVs on each of the branches of the tree to mutational signatures detected in an external cohort of MPNSTs, revealing the platinum-associated signature SBS35 in all branches but not the MRCA. Otherwise, mutational processes remained stable during the evolution of this tumour (**Supplementary Fig. 4**).

To further explore the heterogeneity of this tumour, we performed single-cell DNA sequencing (scDNAseq) using 10x Genomics CNVkit and Direct Library Preparation^7,20^ (DLP+) on the same samples at a median depth of 0.02X per cell. To corroborate the mutation clusters inferred from our bulk subclonal reconstruction, we genotyped variants in the single cells and assessed their co-occurrence patterns. Validating our phylogenetic tree, co-occurrence was only seen for pairs of SNVs either in the same subclonal mutation cluster or across clusters that were part of the same lineage (**Supplementary Fig. 5**).

We derived integer copy-number profiles for 6,039 tumour cells (**Methods**, **Supplementary Fig. 6**). This revealed complex intra-regional heterogeneity with multiple subclones distinguishable by their CNAs in each region. We utilised these single-cell copy-number profiles to define 18 subclones using a K-means clustering approach (**Methods**). The majority of subclones were profiled by both technologies (**Fig. 1d-e**). However, the P_1 and P_2, and P_3 and R4_1 subclones were only profiled with 10x and DLP, respectively, likely due to sampling differences. Dimensionality reduction of all CNA profiles confirmed that the CNA profiles of most single cells clustered with those of their corresponding bulk regions (**Supplementary Fig. 7**). To identify additional subclone-specific SNVs, we performed *de novo* somatic variant calling on the 10x scDNAseq data (the analogous analysis on the DLP+ data was noisier; **Methods**). To account for the higher levels of technical noise in single-cell data, we trained a machine learning classifier on a manually annotated subset of candidates to filter and identify a further 1,107 SNVs. By genotyping cells in the same subclone for SNVs and performing hierarchical clustering, we were able to augment the SNV clusters derived from bulk sequencing and identify SNVs specific for subclones originating from the same region (**Supplementary Fig. 8**).

The near genome-wide LOH observed in this tumour allowed us to reconstruct chromosome-scale haplotypes from the bulk tumour data. Using these phased haplotypes, we fitted single-cell phased allele-specific copy number profiles from LogR and BAF values calculated for genomic bins (**Methods**). These single-cell allele-specific copy-number profiles allowed us to refine the phylogenetic tree of this tumour down to the single-cell level using a CNA-based maximum parsimony approach (**Fig. 1f**).

Taken together, this case of MPNST displayed extreme levels of chromosomal instability: at the bulk level, the entire genome was observed to exhibit either CNAs or copy neutral LOH. Even more strikingly, at the single cell level, 99.5% of the genome had subclonal events (supported by at least 10 cells for each copy-number state), with a median of 3 different allele-specific copy-number states per segment.

### Microdissection and mini-bulk sequencing uncover intertwined local clonal expansions

As a result of the extensive heterogeneity, even neighbouring samples from the same tumour region may comprise different subclones, complicating analyses. This was exemplified by subclone R1_1, displaying a large loss on chromosome 10. While this clone accounted for ∼35% of cells in the R1 single-cell data, it was absent from or contributed little to the bulk sequencing data (**Fig. 1b, e**). Note also that no subclone carrying a reciprocal gain was identified, which could have masked the loss at the bulk level and provided an alternative explanation. As the samples for bulk and single-cell DNA sequencing were taken from adjacent but not identical tissue cuts, this points to the presence of a microscopic subclonal expansion, potentially arising from selection, or due to spatial effects (i.e. clonal intermixing and sampling bias).

To explore this further, we sought to characterise subclones in close spatial proximity to each other in more detail and ascertain their phylogenetic relationships. We performed laser capture microdissection (LCM) from three sections lining the front, side and back of tumour blocks from the primary, R1 and R4 regions (**Fig. 2a**). We captured and successfully amplified DNA from 101 LCM spots and performed mini-bulk whole-genome sequencing with a median coverage per spot of 1.35X. We derived phased allele-specific CNA profiles from these mini-bulk LCM samples (∼300 µm diameter). While coverage was low, these showed little evidence of subclonality within them. However, only one of the nine sections exhibited a correlation between genetic and physical distances (on the order of millimetres) between spots, indicating complex spatial patterns of concurrent clonal expansions with intertwining tumour growth paths but not complete admixing (**Supplementary Fig. 9**). To aid exploration of spatial relationships between subclones, we developed an interactive R Shiny app, available at https://vanloo-lab-mpnst-lcm-shiny.share.connect.posit.cloud. These analyses revealed patterns of spatial copy number evolution such as gradients of segmental copy-number gains (**Fig. 2b**).

**Figure 2.**
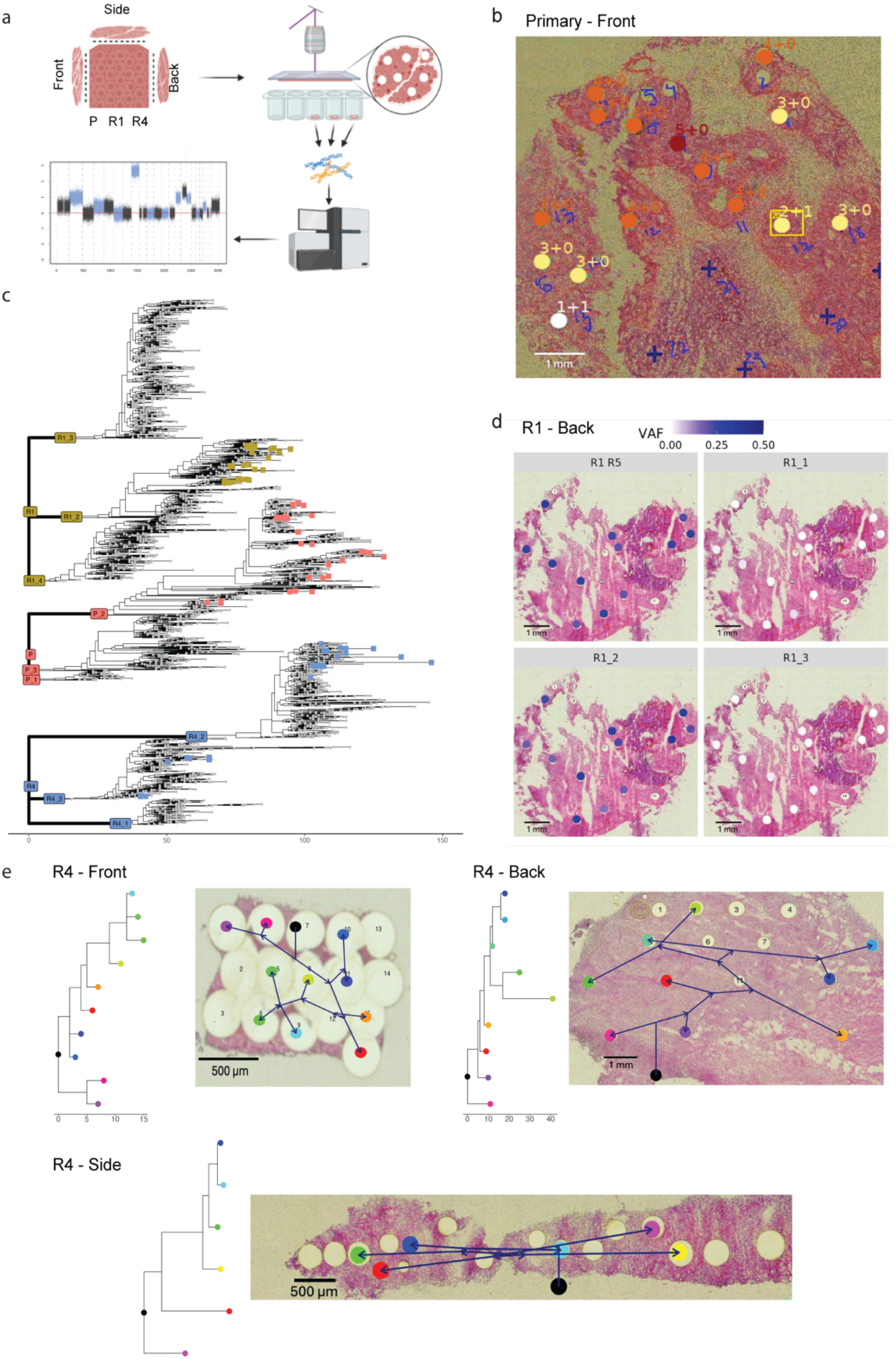
Laser capture microdissection reveals spatial relationships of tumour subclones. (**a**) Experimental design of spatial profiling through LCM. (**b**) Spatial distribution of copy number state in spots shown for chr2:36.4Mb-88.9Mb for the front section of the primary tumour. (**c**) LCM mini-bulk spots shown as squares coloured by their region of origin on the relevant clade of the full resolution phylogenetic tree. (**d**) Variant allele frequency (VAF) of cluster of subclone specific SNVs for the shared clonal cluster between R1 and R5, and subclone specific clusters for R1_1, R1_2 and R1_3 in the back section of R1. (**e**) MEDICC2 phylogenetic relationships of mini-bulk spots projected onto histological images of the front, back and side sections of R4 with arrows showing the relationship between spots. Captured LCM spots are shown in colour and the MRCA is shown in black.

We next utilised the mini-bulk allele-specific copy number profiles to place spots onto the fully refined phylogenetic tree based on the closest single-cell by event-distance, revealing several attributes of the physical expansion in this tumour. Firstly, as expected, mini-bulk spots were placed exclusively with subclones from their corresponding region (**Fig. 2c**). However, they did not always cluster by the section they were captured from, which is in line with the notion of localised clonal expansions and deep intertwining of subclones within cm^3^ volumes (**Supplementary Fig. 10**). Furthermore, spots from R4 were representative of two subclones sampled for the single-cell DNA analysis, whereas those captured from the primary tumour and R1 of the recurrence only derived from one specific subclone (**Fig. 2c**). Genotyping subclonal clusters of SNVs in LCM spots confirmed that only SNVs specific to R1_2, but not other R1 subclones, were present in sections from R1, in line with CNA-based matching (**Fig. 2d**). This also further validates the subclone-specificity of the *de novo* SNVs identified through scDNAseq (**Fig. 1f**). Finally, we reconstructed the temporal acquisition of CNAs of mini-bulk spots using MEDICC2. This enabled us to project the phylogenetic relationship between spots onto spatial locations through simple interpolation (**Methods**), suggesting intricate tumour growth paths at high magnification (**Fig. 2e**).

### Copy number alterations are the main driver of RNA expression differences between single tumour cells

We next performed scRNA sequencing on each of the six tumour samples to profile the cellular composition and assess the impact of copy number heterogeneity on cell transcriptomes. Profiling a total of 37,716 cells, we performed dimensionality reduction, unsupervised clustering and visualisation by UMAP (**Fig. 3a**, **Supplementary Fig. 11**, **Methods**).

**Figure 3.**
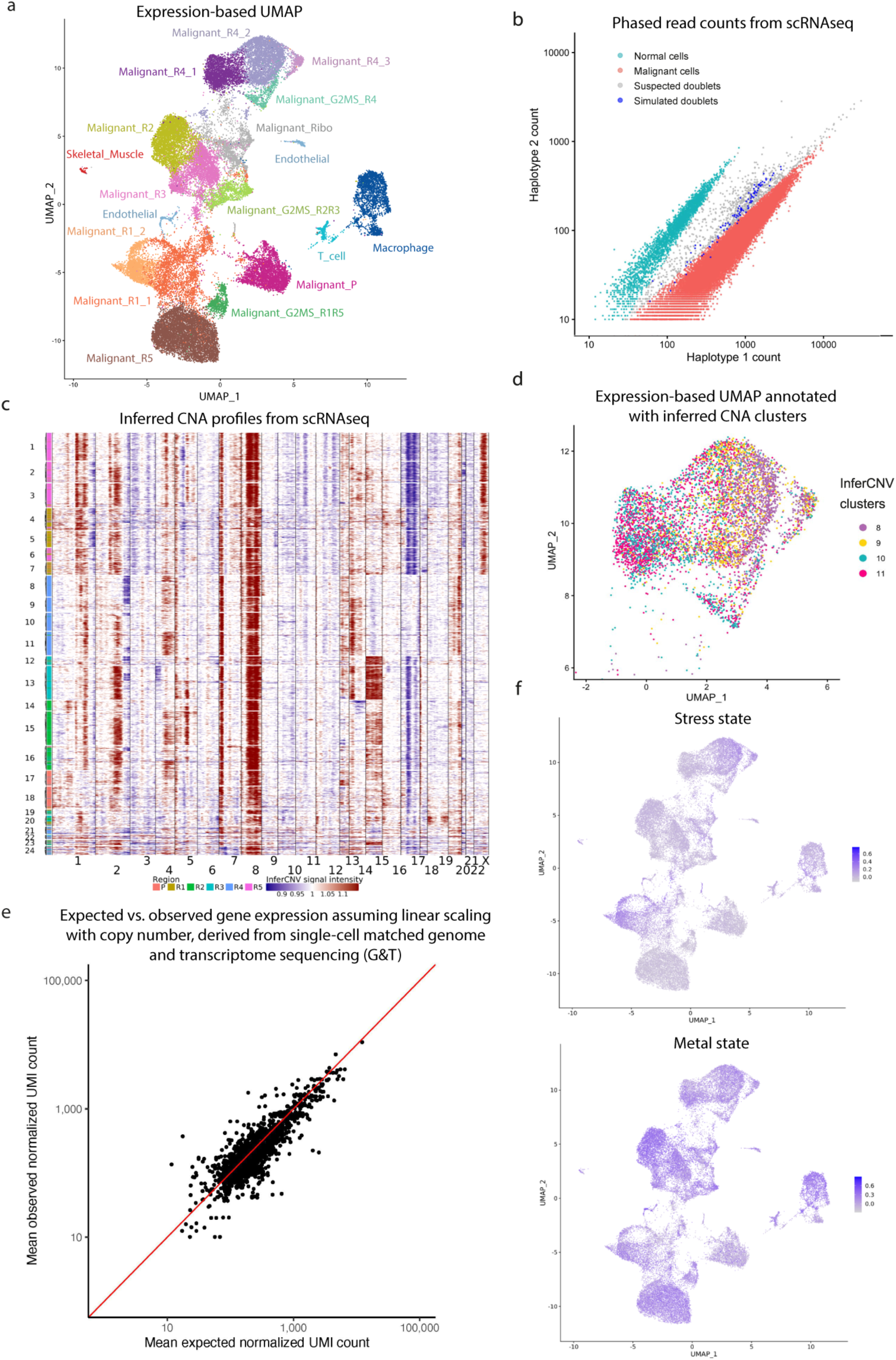
Single-cell gene expression heterogeneity is underpinned by CNAs. (**a**) UMAP of scRNAseq data with cell type annotations. Tumour cell clusters are named by their predominant region of origin and clusters of cycling cells are named CC. (**b**) Total phased scRNAseq read counts per cell from the retained (1) and lost haplotype (2), coloured by their tumour/normal annotation. Simulated doubletsx are in blue, suspected tumour-normal doublets in gray. (**c**) InferCNV copy number profiles with gains and losses relative to normal TME cells shown in red and blue respectively. Cells are clustered using k-means and annotated by their region of origin. (**d**) UMAP of the R4 scRNAseq data only, with cells coloured by their inferCNV-defined copy number cluster. (**e**) Expected vs. observed mean expression of genes in cells with higher copy number states, derived from matched genome and transcriptome sequencing (G&T-seq). Expected values were calculated based on the mean read-depth normalized UMI counts from cells at lower copy number states. (**f**) Activity of the “Stress” and “Metal” meta-programs shown on UMAP.

Marker gene analysis showed that the tumour microenvironment (TME) was composed mostly of macrophages (6.7% of all cells, range: 2.2-10.1%), T cells (1.1%, range: 0.2-2.6%), endothelial cells (1.1%, range: 0.3-2.8%) and skeletal muscle cells (0.3%, range: 0.0-1.0%). The remaining cells (90.9%, range: 83.7-97.1%) clustered by region of origin and were identified as tumour cells. Their high proportion matched the purity estimates of the bulk samples.

To confirm our cancer cell annotations, we leveraged the near genome-wide LOH in this tumour. Using the phased haplotypes, we genotyped single cells at heterozygous SNPs and generated phased B-Allele Frequency (BAF) values representing the minor alleles across the genome. As expected, cells identified as malignant cells had fewer reads reporting the lost alleles than TME cells (**Fig. 3b**). This showed to be a marker gene-independent method for confirming not only cell type, but also to identify potential tumour-normal doublets, whose expression frequently clustered with normal cells (**Supplementary Fig. 12**, **Methods**).

To explore CNA heterogeneity within the scRNAseq data, we inferred CNA profiles using inferCNV (**Fig. 3c**). Cells with similar CNA profiles were in close proximity on the UMAP manifold (**Fig. 3d**, **Supplementary Fig. 13a-b**), suggesting that at least part of the transcriptomic similarities were driven by copy number differences. To quantify this, we calculated the Jensen-Shannon distance, an information distance quantifying how differently cells allocate their transcriptional activity across genes, between transcriptomes from different inferCNV clusters and between cell types (**Methods**). Distances between cells in distinct inferCNV clusters were higher than those within the same cluster (including within the normal), but intermediate compared to distances between cells of different types (**Supplementary Fig. 13c)**, indicating that ongoing CNA evolution substantially reshapes transcriptomes across tumour subclones, though to a lesser extent than the reshaping of trans-differentiations between cell types.

Gene dosage effects have been detected across tumour types through analysis of bulk data^21^. To rigorously quantify this at the single-cell level, we performed simultaneous genome and transcriptome profiling of 158 single cells using G&T-seq, and derived copy number profiles and gene expression counts. Linear gene dosage relationships were observed for nearly all genes located on segments with differing copy number states across tumour cells (**Fig. 3e**, **Supplementary Fig. 14**). This confirms that CNA-based gene dosage effects shape phenotypic diversity, and explains why cells within the same CNA clusters have more similar transcriptomes.

Copy-number changes co-modulate the dosage of genes in-*cis*. However, it is unclear how CNAs impact expression in-*trans*, through gene regulatory network activity, which we quantify using SCENIC^22^, or how they affect the activity of previously identified “meta-programs” shared across many cell systems^23^. First, we observed that tumour cells with similar CNA profiles were in close proximity in the gene regulatory network UMAP space (**Supplementary Fig. 15a-b**). Quantification using pairwise Adjusted Rand Index (ARI) confirmed the correspondence between CNA and gene regulatory network clusters, however CNA clusters were typically spanning at least two gene regulatory network clusters, indicating that CNA heterogeneity only partially shapes the overall regulatory network landscape (**Supplementary Fig. 15c**). Notably, only two gene regulatory network clusters corresponded to cell cycle phases and were characterized by a FOXM1 regulon, while the other regulons were related to processes such as immune response, metabolism, or editing. Second, we scored each individual cell for the previously identified meta-programs and noted differences between and within regions. Specifically, cells from all five recurrence regions had a higher “Metal” cell state than those from the untreated primary, in keeping with adaptation following systemic anti-cancer therapy (**Fig. 3f**). Furthermore, we noted increased states of “Stress” in a subset of cells from R1 and R4 (**Fig. 3f**, **Supplementary Fig. 16**). These findings highlight the ability of those meta-programs to identify shared cell states of genetic subclones, despite their globally different transcriptomes.

### Spatial profiling reveals genetic and spatial relationships between tumour subclones

Next, to map transcriptomic effects in a spatial manner and profile the makeup of this tumour, we performed 10x Visium spatial transcriptomics on 8 slides (one from R1, R2, R3, R4, and two each from the primary tumour P and R5). We genotyped heterozygous SNPs to determine the malignant/non-malignant composition of each spot. Virtually all spots had a constant ratio of counts of each haplotype, which was similar to those seen for tumour cells in the scRNAseq data (**Supplementary Fig. 17a**). As spots typically cover several cells, the transcriptomes of all spots were likely composed of mostly tumour cells and the occasional TME cell. However, the malignant cell fraction could be overestimated, since they may have higher transcriptional output^24^ (**Supplementary Fig. 17b**). Therefore, we performed deconvolution of the spots with CARD, a method that uses matched scRNAseq to extract the local expression contribution of different cell types^25^. This confirmed that most spots contained a mixture of malignant and non-malignant cells and identified distinct regions of macrophage infiltration in the primary tumour (**Fig. 4a**). In contrast, the TME was relatively uniform in all other samples. This mixture of TME and tumour cells in most regions was confirmed following histological review of haematoxylin and eosin (H&E) stained slides.

**Figure 4.**
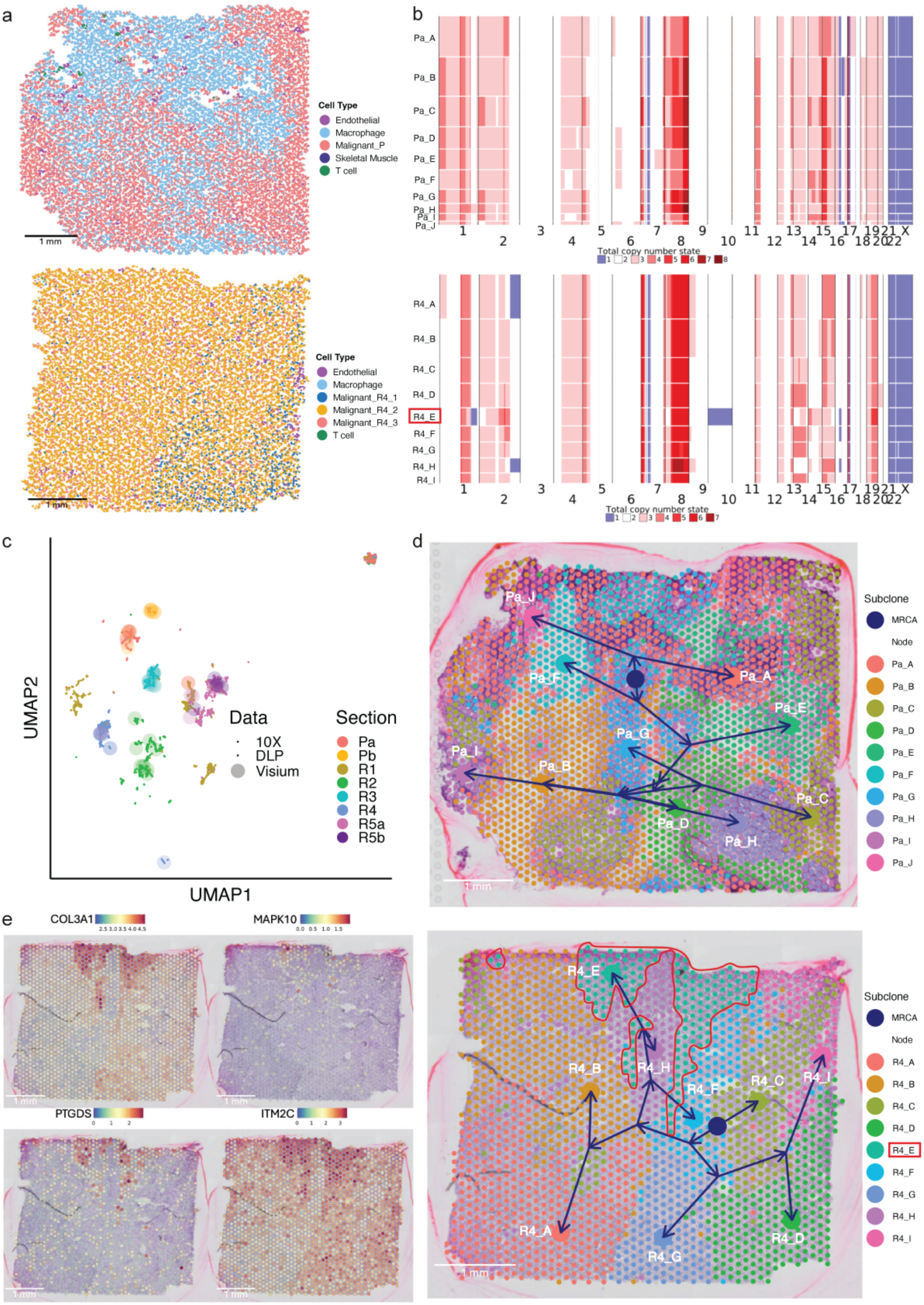
Spatial transcriptomics allows *in situ* phylogenetic reconstruction of tumour subclones. (**a**) CARD single-cell deconvolved 10x Visium spatial transcriptomics sections of the primary (Pa) and R4 sections. (**b**) Inferred integer copy-number profiles of clusters for Pa and R4. R4_E cluster annotated in red. (**c**) Copy number profiles of Visium clusters projected onto scDNAseq UMAP. (**d**) Spatial phylogenetic trees superimposed on Pa and R4 slides. Spots are coloured by their expression-based cluster and centroids of Visium clusters were used to position terminal nodes with the MRCA shown in dark blue. Spots belonging to the R4_E cluster are outlined in red. (**e**) Top upregulated genes in R4_E cluster by differential expression analysis comparing R4_E with all other spots.

CNAs may also be inferred from spatial transcriptomics data^9^. Since no representative normal Schwann cells were profiled and the relationship between gene expression and DNA copy number was predominantly linear (**Fig. 3e**), we used the R1 slide as a copy-number homogeneous reference to detect CNAs in the other sections (**Supplementary Fig. 18a,b**). Leveraging the consensus single-cell copy-number profile of an R1 subclone, relative changes were used to calculate total copy-number profiles from the gene expression-based ratios, revealing subclonal heterogeneity in most sections (**Fig. 4b**, **Supplementary Fig. 19**). Supporting the accuracy of this approach, the inferred subclonal profiles aligned with those detected through scDNAseq (**Fig. 4c**). Indeed, subclones present on tissue sections Pa and Pb matched those of the P_2 and P_3 clusters in the scDNAseq, respectively, revealing that the spatial separation of distinct transcriptomic clusters is driven by their underlying genetic relationship (**Fig. 1e**, **Supplementary Fig. 19**). Furthermore, the R4_E cluster of spots displayed losses on chromosome arm 1q and chromosome 10, which were consistent with the R4_1, and markedly different from other clusters on the tissue section (**Fig. 1e**, **Fig. 4b**, **Supplementary Fig. 18b**). Spatial phylogenetic trees for the centroids of each spatial subclone were reconstructed and superimposed upon each tissue section (**Fig. 4d**). Although these could not be directly compared, they were reminiscent of those seen in the LCM data. Genes differentially expressed between R4_E and other R4 subclones showed no clear pattern of CNAs (**Fig. 4**, **Supplementary Fig. 20**). R4_E upregulated genes included several involved in neuronal development or signalling such as *COL3A1*, *MAPK10*, *PTGDS* and *ITM2C*, hinting at co-opting of these signalling pathways. Overall, these analyses provide a high-resolution view of tumour growth dynamics and demonstrate that spatial gene expression patterns closely recapitulate the genetic relationships between subclones in MPNST. At the same time, trans-acting influences, including meta-programs and regulons, remain key drivers of transcriptional heterogeneity.

## Discussion

In this study, we combine state-of-the-art bulk, single-cell and spatial sequencing approaches to meticulously characterise the spatiotemporal evolution of an MPNST, a rare sarcoma. This allowed us to reconstruct its evolutionary history with unprecedented granularity.

The primary tumour was genomically highly unstable and had undergone near haploidization of its genome. This phenomenon has been observed in sarcomas and other tumour types, although the underlying mechanism as well as its clinical significance remains unknown^26^. Recent work on a large cohort of MPNSTs has identified this as a critical step in MPNST tumorigenesis^27^. We observed multifurcation events at multiple levels, resulting in a branching phylogenetic tree. This initial multifurcation could be explained by polyphyletic seeding of tumour cells prior to chemotherapy that survived due to dormancy, but then escaped dormancy and rapidly proliferated to give rise to multifocal recurrence. Ongoing chromosomal instability, generating detectable subclones with distinct CNA profiles, likely contributed to subsequent branching.

We leveraged this CNA-based native barcoding system for several purposes. Firstly, we identified clusters of subclones harbouring specific SNVs to increase the resolution of the phylogenetic tree. Furthermore, we used CNA profiles from four different modalities (10x and DLP+ scDNAseq; LCM mini-bulk and bulk WGS) to generate a common minimum event distance-based phylogenetic tree, revealing spatial patterns of tumour growth. Finally, by leveraging G&T-seq, we demonstrated that CNAs are the major determinant of gene expression heterogeneity in these MPNST cells.

Despite recent studies investigating how different clones compete and expand^28,29^, our knowledge of how tumours grow in 3D remains limited. This is in part due to difficulties in determining the phylogenetic relationship between cells in a spatial context. By applying LCM, we identified local expansions and, to our knowledge, for the first time traced subclonal lineages at scale in a human tumour. Furthermore, we characterised the interaction between tumour cells and the TME, and extend lineage tracing into spatial transcriptomics. We also showed that spatial RNA-based copy number inference, when constructed carefully and in a tumour with large-scale CNAs, can reliably reproduce DNA-based profiles. This highlights the added power of spatial transcriptomics approaches to start studying genotype-to-phenotype relationships in chromosomally unstable tumours.

Overall, this MPNST and its multifocal recurrence offer a glimpse into the degree of evolutionary detail that can be extracted by utilising native barcodes. As maturing technologies such as Primary Template-directed Amplification (PTA) continue to increase the efficiency and amount of information captured from a single-cell, we anticipate studies using *in vivo* lineage tracing will reveal ever more fine-grained details. Our current multimodal analyses required utilising and optimising a wide suite of existing packages, benefitting future efforts. Notably, we expanded upon a current CNA-based phylogenetic tree reconstruction method to perform phylogeny reconstruction and lineage tracing using allele-specific CNAs in single cells^19^. In addition, we developed and validated custom approaches such as *de novo* SNV calling and filtering from scDNA-seq and detection of tumour-normal doublets for single-cell and spatial transcriptomics data.

By focusing on exceedingly in-depth characterization of an individual tumour, we highlight the level of detail current technologies provide to resolve phylogenies. Our results offer a glimpse of a complete spatio-temporal representation of tumour development, linking genotype to (molecular) phenotypes. Nevertheless, while genotypes and phenotypes can be observed in 3D space, temporal inference from a snapshot in time is only possible for genotypes. Time series analyses using cancer models could help fill this gap and complement *in vivo* observations and inference. When applied to further datasets, integration of these modalities will be pivotal in uncovering the ancestry and phenotype of clinically relevant subclones. Multi-omics data integration studies such as ours may shed light on the critical events in cancer development and progression, and potentially lead to methodical subclone-informed treatment strategies.

## Methods

### Patient sample acquisition

The primary tumour was collected from surgery at the Royal National Orthopaedic Hospital and snap frozen in liquid nitrogen. A total of 5 local recurrence region samples were collected from amputation tissue and snap frozen in liquid nitrogen or collected in PBS, manually minced and dissociated into single-cell suspensions using Collagenase II. All samples were then transferred to −80°C for long-term storage. Clinical data were collected by the Royal National Orthopaedic Hospital biobank, London Sarcoma Service databases and pathology archives. The individual gave their written and informed consent to provide samples for this study, which was approved by the National Research Ethics Service (NRES) Committee Yorkshire and The Humber, Leeds East (15/YH/0311).

### Bulk sequencing

DNA was extracted from germline blood and tissue sections cut from tumour samples. Libraries were made from native DNA using the NEBNext Ultra II FS DNA Library Prep kit (New England BioLabs) and 100 base paired-end sequencing was performed on the HiSeq 4000 system according to Illumina protocols. Paired-end reads were aligned to the reference human genome (GRCh38) with BWA-MEM^30^. Duplicates were marked using MarkDuplicates (Picard v2.19.1) and base quality scores were recalibrated with Genome Analysis Toolkit (GATK v4.1.2.0) BaseRecalibrator^31^.

### Single-cell sequencing

Nuclei were isolated from all tumour samples using EZ Prep (Sigma-Aldrich) and used as input for single-cell barcoding using the 10x Genomics Chromium platform and DLP+^20^. For scDNAseq, the Chromium Single Cell CNV kit (10x Genomics) was used and for scRNAseq, the Chromium Single Cell 3’ v3 kit (10x Genomics) was used. The 10x Genomics-generated DNA/cDNA libraries were sequenced on an Illumina HiSeq 4000 using 150 base paired-end and 100 base single-end sequencing, respectively. Reads for scDNAseq were aligned to GRCh38 using BWA-MEM (v0.7.17)^30^. Cellranger-DNA (v1.1.0) and Cellranger (v3.0.1) were used to demultiplex and align reads to GRCh38 for scDNAseq and scRNAseq, respectively^32^. In total, 37,716 scRNAseq cells passed the Unique Molecular Identifier (UMI) noise filter. A total of 6,795 10x scDNA-seq cells passed the initial 10x barcode cell-calling filter; after ASCAT.sc quality filtering and integration with DLP+ single-cell DNA profiles, the final copy-number dataset comprised 7,426 cells, including 4,426 10x-derived and 3,000 DLP+-derived cells. For DLP+, nuclei were stained with Hoechst and 1,225 nuclei per sample were spotted into 5,184 nanowell chips preprinted with unique dual index barcodes using a CellenOne cell sorter^20^. Pooled barcoded single-cell libraries were sequenced on an Illumina Novaseq using 150 base paired-end sequencing.

### Flow cytometry-based ploidy analysis

Small fragments of tumour tissue were placed in Pepsin-HCl solution (5 mg/ml pepsin in 10 mM HCl solution). A Bioruptor sonication system (Diagenode) was used to dissociate the tissue and incubated at 37°C for 30 minutes. The solution was then spun down and washed with 1X PBS. 1 ml of 70% ethanol was added drop-wise whilst vortexing, then incubated at room temperature (RT) for 1 hour before a further spin and wash with 1X PBS. The sample was then incubated with propidium iodide (1 µg/ml) overnight at 37°C. Flow cytometry was performed on a LSR II Flow Cytometer (BD Biosciences). Nuclei were gated based on the FSC vs. SSC plot, then single cells were gated on the area vs. height plot for the yellow-green laser with 610/20 bandpass filter. Histograms were plotted for each region, revealing subpopulations with different ploidies. The ploidy for each peak was calculated by taking its median signal intensity value and dividing by the median of the diploid peak then multiplying by 2.

### Laser capture microdissection

Tumour tissues from the primary tumour, R1 and R4 were embedded in optimal cutting temperature (OCT) compound and stored at −80°C. Sections 16 μm thick were cut from the front, side and back of tissue blocks, mounted on PEN-Membrane Slides (Leica) and stained with H&E. LCM of 16 to 24 spots per section was performed according to the manufacturer’s protocol on the Leica LMD7000 system. Whole-genome amplification was performed on extracted DNA using the Genome Plex kit (Sigma-Aldrich). Libraries were prepared from 134 successful amplifications using the NEBNext Ultra II FS DNA Library Prep kit (New England BioLabs) and 100 base paired-end sequencing was performed on an Illumina HiSeq 4000. Reads were aligned to GRCh38 using BWA-MEM (v0.7.17)^30^.

### Genome and transcriptome sequencing

Single cells dissociated at time of tissue collection were sorted into 96-well PCR plates for the recurrence regions. Nuclei were prepared from the primary tumour and also sorted into a 96-well PCR plate for G&T-seq. Unfortunately, scDNAseq for the R1 region failed. In total, 216 single-cell genomes and 300 single-cell transcriptomes were successfully sequenced resulting in 158 paired genomes and transcriptomes from the same cell. Genomes were aligned to GRCh38 BWA-MEM (v0.7.17)^30^ and duplicates were marked with SAMtools markdup (v1.3.1)^33^. Transcriptomes were aligned to GRCh38 and gene counts generated with STARsolo (v 2.7.1a)^34^.

### Spatial transcriptomics

All tumour tissues were embedded in OCT and stored at −80°C. 10 μm sections cut for each tissue and processed using the Visium Spatial Gene Expression Kit (10x Genomics). Tissues were permeabilised for 3 minutes after optimisation of permeabilisation conditions. Sections were stained with H&E and imaged using a VS120 Slide Scanner (Olympus Life Science) under 20x magnification. Visium libraries were constructed according to 10x protocols and sequenced on an Illumina HiSeq 4000. SpaceRanger (v1.1.0; 10x Genomics) was used to demultiplex reads and align to GRCh38 after manual alignment of fiducial markers and marking of tissue boundaries for each slide using Loupe Browser (v4.1.0; 10x Genomics).

### Copy number and variant calling in bulk sequencing data

Battenberg (v2.2.9) was used to obtain subclonal copy-number profiles for the primary tumour and each region of the recurrence^17^. The algorithm first counts the number of reads for each allele at SNP positions from the 1000 Genomes Project in the tumour and normal BAMs. The counts from SNPs are then used to derive a logR (log2 of the normalised ratio between tumour and normal reads) and BAF (ratio between the paternal and maternal alleles). Battenberg then uses population-based phasing for haplotype reconstruction with Beagle5^35^. Segmentation by Piecewise Constant Fitting (PCF) is then performed on the logR and BAF values of heterozygous SNPs to determine genomic segments with the same copy-number state^36^. Tumour purity and clonal copy number states are co-estimated using logR and BAF values and a one-sided t-test is performed on the BAF values against the expected BAF for clonal copy- number, to determine if there is evidence of subclonal copy number event for a genomic segment - the null hypothesis is rejected if the t-test p-value<0.05.

As several samples were available, the multi-sample mode was used with external phasing information for haplotypes also incorporated from linked-read sequencing performed on the same tumour samples to improve the phased haplotype blocks and segmentation. Most tumour samples were highly pure, resulting in BAF values very close to 0 or 1 which, given the finite tumour sequencing coverage, confounds detection of subtle shifts in BAF. Therefore, to artificially reduce the purity, *in-silico* spike-in of reads from the matched normal was performed, which resulted in improved segmentation and copy-number calling. The modified Battenberg plot to display copy number profiles of all regions was created by overlaying the CNA profiles of all samples and annotating the total copy number state by region.

Somatic variants were called using Mutect2 (GATK v4.1.8.0) in a Singularity container (v3.6.4)^31,37^. Variant Call Format (VCF) files were then merged and indexed and MergeMutectStats from GATK was used to combine the stats files for each chromosome. The LearnReadOrientationModel, GetPileupSummaries and CalculateContamination steps in the GATK pipeline were then run to allow SNV filtering using FilterMutectCalls and those which passed were annotated with Funcotator^31^.

### Mutational clustering and phylogenetic tree reconstruction from bulk sequencing data

Multidimensional Bayesian Dirichlet Process-based mutation clustering (ndDPClust) (v2.2.8) was run in multi-sample mode on all SNVs present in all regions^18^. This algorithm identifies clusters of mutations belonging to subclones based on their CCFs. The R package dpclust3p (v1.0.8) was used to prepare inputs for ndDPClust. Only SNVs on chr1-22 and X were retained. Due to memory constraints, all combinations of 4 from 6 samples were run for 12,000 iterations (2,000 burn-in). ClusterID-based consensus clustering (CICC) was used to generate 40 consensus clusters across the 15 runs and allowed calculation of CCFs of each cluster^1^. This revealed clonal clusters for all regions and a subclonal cluster for R4. Subclonal clusters of SNVs for other regions were identified from a large cluster of SNVs with low CCF (∼0.05) by extracting SNVs with CCF of 0 in all other regions. Remaining clusters which violated the pigeon-hole principle or crossing rule such as those with constant CCFs across samples were removed due to likely being artifactual clusters. Based on these criteria, 2,223 out of the total 13,957 SNVs were filtered out. The remaining clusters were taken forward for further analysis. The phylogenetic tree was reconstructed manually by assessing the CCFs of each subclone in each region. The pigeon-hole principle was then applied to mutational clusters only present in one region to determine their relationship (i.e. branching vs. linear). This principle states that if the CCFs of two mutational clusters sum up to more than that of their shared ancestral cluster, they must be collinear.

### Mutational signature analysis

The R package Sigfit (v2.2.0) was used to fit SNVs from each mutational cluster against a catalogue of mutational signatures found in a cohort of MPNSTs from Genomics England (GEL)^38^. These included signatures SBS1, SBS2, SBS5, SBS8, SBS9, SBS13, SBS17a, SBS17b, SBS18, SBS28, SBS30, SBS35, SBS36 and SBS39 (COSMIC v3.2)^39^. Sigfit^38^ was run for 10,000 iterations with a burn-in of 5,000 iterations with each mutational cluster as a sample, thereby deriving the mutational signatures for each clonal/subclonal cluster. Mutational signatures with a 95% highest posterior density interval lower limit of greater than 1% of SNVs in any cluster were retained (signatures SBS13, SBS17a, SBS17b, SBS28 and SBS36 were removed) and mutations were refitted to the remaining signatures.

### Total and allele-specific copy number calling in single cells, LCM and G&T-seq

ASCAT.sc (v0.1) was used to infer copy number profiles from the combined 10x and DLP+ scDNAseq data (available at https://github.com/VanLoo-lab/ASCAT.sc). First, pre-computed 30kb bins for GRCh38 included in ASCAT.sc were loaded. These bins are selected to have the most stable counts in a panel of diploid normals. Read counts were then derived in each bin with MAPping Quality (MAPQ)>=30 excluding duplicates for each 10x or DLP+ sample. Log read counts for each group of consecutive 30kb bins (leading to 480kb genomic bins) were then smoothed by applying a loess fit against GC-content to obtain the corrected logR.

To identify purity and ploidy in a similar fashion to ASCAT^40^, a grid search was then performed on all ploidy values between 1.7 and 5 by steps of 0.01 and a purity of 0.5 or 1 to fit copy number profiles from the logR track. Noisy cells were filtered out based on the average standard deviation of the logR across segments and number of reads per bin per tumour chromosomal copy (NRPCC). Cells passing this filter from all samples were then combined and multi-PCF (mPCF) was then used for segmentation of shared genomic segments with different copy number states across all cells. This resulted in cells with satisfactory copy number profiles with shared breakpoints. Fitted integer copy number profiles revealed populations of cells present in all regions which appeared to have undergone further whole-genome doubling (WGD). Whilst these sub-populations were confirmed on FACS ploidy analysis, they were removed from further analysis as it was not possible to accurately assign which cells had undergone WGD.

A K-means approach with K set to 22 clusters was then applied to the remaining cells to assign cells to copy-number based subclones. Clusters with fewer than 5 cells, a cluster of cells with noisy profiles and a cluster of normal cells were removed. This yielded 18 tumour subclones with 6039 cells in total which were used in subsequent analyses. A UMAP dimensionality reduction of all single-cell copy number profiles using Manhattan distances was performed with the R package umap (v0.2.7). Copy number profiles of bulk samples were then transformed or projected into this learned space. Similarly, copy number profiles inferred from Visium spatial transcriptomics were also projected onto this shared UMAP.

Allele specific copy-number profiles were then derived using ASCAT.sc by leveraging BAF values calculated from the phased haplotypes derived from the bulk Battenberg runs. As single-cell data is particularly sparse, logR and BAF are clustered across cells to improve their estimation and thus, allele-specific copy number profile fitting. Copy number profiles were plotted by taking the integer value for each genomic bin per cell and plotting them as a heatmap using a custom wrapper function around the ComplexHeatmap package which annotates the chromosomes. For allele specific copy number profiles, the states of the two alleles were converted to a single value to allow plotting as a heatmap in a similar fashion.

ASCAT.sc was also used to perform integer and allele-specific copy number calling on shallow coverage WGS data from LCM spots and G&T-seq cells. This was done by applying the breakpoints from the scDNAseq data to the LCM and G&T-seq logR and BAF tracks to create recurrent breakpoints across sequencing methods and allowing for ploidies between 1.7 and 5.

### Genotyping SNVs in single cells

Clusters of mutations identified in bulk WGS through DPClust^18^ and CICC^1^ were genotyped in the scDNAseq data using alleleCount (v4.0.0; https://github.com/cancerit/alleleCount). A binary co-occurrence matrix was generated where SNVs were considered co-occurring if their ALT alleles were supported by at least one read in the same single cell.

### De novo SNV calling and subclonal mutation clustering in 10x scDNAseq data

Somatic variant calling was performed using the 10x scDNAseq data. First, mutations were called using Mutect2 against the bulk normal as well as pooled scDNAseq normal cells as matched controls^31,37^. scDNAseq cells were identified as normal by having diploid copy number profiles and a 1:1 ratio of the two haplotype counts. Only shared SNVs detected against both bulk and pooled normal were retained. Next, a XGBoost machine learning classifier from the R package xgboost (v1.3.2.1)^41^ was trained to filter out false SNV calls. The following features of SNVs were used for statistical modelling and as input for the XGBoost algorithm: number of reads with the reference and alternative allele in the bulk tumour, number of reads with the reference and alternative allele in pooled scDNAseq tumour cells, number of reads with the reference and alternative allele in the bulk normal, number of reads with the reference and alternative allele in pooled scDNAseq normal cells and whether the SNV is within 1Mb of the centromere. A manually curated subset of 200 SNVs was used to train the classifier reaching 96% accuracy on a hold-out set of 100 SNVs after 100 iterations.

Clusters of SNVs specific to subclones were determined by pooling cells for each subclone together and genotyping SNVs. SNVs were considered present in a subclone if one read with the alternative allele was found in at least 3 cells from the subclone. Clustering of subclone specific SNVs was performed using hierarchical clustering (*hclust* R function with the ward.D2 method) after removal of SNVs assigned to be truncal by ndDPClust and SNVs present on 1000 Genomes Project SNP positions.

### Single-cell phylogenetic tree reconstruction

The SNV-based subclone tree was used for reconstructing the main branches of the phylogenetic tree. Single cells from each subclone were then used to generate a minimum-event distance tree with MEDICC2 (v0.5b) with WGD events enabled and minimum segment length of 1.5Mb^19^. These single-cell trees were then grafted onto subclone tips to create a phylogenetic tree refined down to the single-cell level.

### Analysis of LCM spots

MEDICC2^19^ was run on the combined scDNAseq and LCM copy number profiles to generate an event-distance matrix. Each LCM spot was then attached to its nearest single cell based on event distance. This phylogenetic tree was then pruned for the regions studied with LCM (primary tumour, R1 and R4) to show the location of LCM spots on phylogenetic trees for these specific regions. The R package Shiny (v1.6.0) was used to create an interactive Shiny app with various features. These include selection of LCM spots to view its allele-specific copy number profile and ratio of haplotype counts, selection of a genomic segment to view the copy-number state in spots for all sides belonging to that region and display of the fully resolved single-cell phylogenetic tree with LCM spots.

MEDICC2^19^ was also used to infer phylogenetic relationships between spots on each individual slide with normal spots excluded. These phylogenetic trees were then represented on histological images by placing successive ancestral nodes equidistant between descendants until the root (MRCA) of the tree was placed. These intermediate node positions were then iteratively adjusted to move 30% of the distance between its original position and its immediate ancestor. Subclone specific SNVs derived from single cells were genotyped in LCM spots from each region as for single-cells to derive the VAF for each cluster of SNVs. These VAFs were then plotted onto the histological image of each slide to show the spatial distribution of SNVs. Manhattan distances of copy number profiles and Euclidean distances from images were used as genetic and physical distance respectively for correlation analysis.

### scRNAseq clustering and cell-type classification

Gene count matrices were loaded into the R package Seurat (v3.2.1)^42^ with removal of cells with more than 5% mitochondrial reads. Counts were then transformed and normalised with the SCTransform function using the top 3000 variable genes. Sample batch correction was not performed as we noted that batch correction was inappropriately removing biological signal. For example, when batch correction was performed using Seurat’s FindIntegrationAnchors and IntegrateData functions, copy-number driven differences between regions were removed with cells from all regions placed together on the UMAP (**Supplementary Fig. 11**). PCA and UMAP dimensionality reduction was performed with 30 principal components used for UMAP calculations. Cycling cells were identified using the function CellCycleScoring with a list of cell cycle markers provided in Seurat.

Clusters were defined using the FindClusters function with a resolution of 0.5. Marker genes for the resulting 17 cell clusters were identified using the function FindAllMarkers. They were then annotated to known cell types based on canonical markers: macrophages (*MRC1*, *F13A1*, *CD163*), T cells (*PTPRC*, *THEMIS*), endothelial cells (*VWF*, *PTPRB*) and skeletal muscle cells (*TTN*, *MYOG*).

The remaining cells were annotated as tumour cells and confirmed to be malignant using a genotyping approach by generating BAF values at 1000 Genomes Project heterozygous SNP positions using alleleCount (v4.0.0) and assigning them to the two haplotypes determined through linked reads of the bulk tumour. Cells annotated as tumour featured significantly fewer reads from the lost alleles compared to TME cells without allelic imbalance. This approach also revealed an additional population of cells with intermediate BAF but annotated as non-malignant cells on the UMAP (**Supplementary Fig. 12a-b**). The BAF of these cells was between the BAF of tumour and normal cells, and exactly matched the expected BAF of simulated tumour–normal doublets which was performed by merging the counts of tumour cells and normal cells (**Fig. 3b**). This genotyping process was also performed to identify cells or spots as normal *vs*. tumour in scDNAseq, G&T-seq, LCM, spatial transcriptomics and Slide-seq data.

### Copy-number alteration calling from scRNAseq

CNAs in single-cell RNA cells were detected using the R package inferCNV (v1.0.2) which uses gene expression intensity across a sliding window of 101 genes to infer copy-number changes^43^. Doublets and cells annotated as normal but with a haplotype count ratio that of tumour cells were excluded. Cells annotated as TME were used as reference normal cells with a cut-off for the minimum average number of reads per gene for reference cells of 0.1, analysis_mode set to “samples” and cluster_by_groups set to true. Chromosome X was also included by setting chr_exclude to “c(“chrY”, “chrM”)”. Hierarchical clustering of correlation distances between inferCNV intensity values using Ward’s minimum variance method yielded 24 clusters.

### Quantification of transcriptomic divergence using Jensen–Shannon distance

To quantify transcriptomic differences between tumour cell clusters and canonical cell types, we computed pairwise Jensen–Shannon (JS) distance across groups defined by copy number and cell identity. Tumour cells were grouped into 24 clusters based on inferred copy number profiles (inferCNV), and normal cells were grouped into four canonical cell types (e.g., T cells, macrophages, endothelial cells, and skeletal muscle).

To mitigate the effect of unequal cell numbers and sampling bias, we generated repeated pseudobulk profiles by randomly sampling N = 100 cells without replacement from each group. For each sampled set, gene expression values were averaged across cells to obtain a pseudobulk profile. This procedure was repeated 10 times per group, resulting in multiple pseudobulk representations for each tumor cluster and normal cell type. All genes were included in the analysis to capture global transcriptomic differences. For each pseudobulk profile, gene expression values were converted into probability distributions by adding a small pseudocount (ε = 1×10⁻¹²) and normalizing across genes.

Pairwise transcriptomic divergence between pseudobulk repeats was quantified using Jensen–Shannon divergence (*JSD*), defined as:

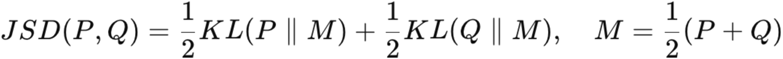

where *P* and *Q* are the normalized gene expression distributions of two groups, and *KL* denotes the Kullback–Leibler divergence. For downstream analyses and visualization, we used the Jensen–Shannon distance (*dJS*), which satisfies the properties of a metric.

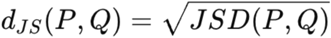

5 types of between-group comparisons were performed using pseudobulk profiles across repeats:

- Tumour vs tumour: between different inferCNV-defined tumour clusters
- Tumour vs normal: between tumour clusters and normal cell types
- Normal vs normal: between different annotated normal cell types
- Within Normal: 10 repeats comparison within one cell type
- Within Tumour: 10 repeats comparison within one tumour cluster

Hierarchical clustering was performed on the Jensen–Shannon distance matrix to assess relationships between tumour and normal groups.

### Gene dosage analysis

Tumour cells profiled using G&T-seq were used to explore gene dosage effects. ASCAT.sc profiles and normalised counts using the SCTransform function were used to obtain the expression and copy number state of each gene in each cell. Only highly expressed genes with expression (SCTransform-based depth-normalized UMI count) of greater than 10 were further analysed. Because zero inflation and dropout in G&T-seq data can cause median expression values to collapse to zero, the mean expression of cells with a lower copy number state which was present in at least 10 cells was calculated. This baseline expression was used to calculate an expected expression for each pairwise comparison with a higher copy number state assuming a linear relationship of gene dosage.

### Gene regulatory network analysis

Gene regulatory networks were identified using the tool SCENIC by running SCENICprotocol (v0.25.0) in a Singularity container^44^. The outputs were then analysed in R with functions from the SCENIC package (v1.1.2)^22^. Dimension reduction of the regulons was carried out using the python package umap-learn (v0.5.1). The top transcription factors for each cluster were identified using the calcRSS function from SCENIC. InferCNV clusters were mapped to the gene regulatory network-based UMAP.

We compared SCENIC clusters and inferCNV clusters using a pairwise adjusted Rand index (ARI). For each pair of clusters, cells were binarized into “in-cluster” and “out-of-cluster” groups for both clustering assignments, and the ARI was calculated between these two binary partitions using the mclust R package. Regulon specificity for each cluster was assessed using the Regulon Specificity Score (RSS). For each cluster, the regulon with the highest RSS was selected as the representative and annotated next to the SCENIC clusters.

### Hallmarks of transcriptional intra-tumour heterogeneity

Genes for all meta-programs from Gavish *et al.* were used to score cancer cell states in individual cells^23^. The AddModuleScore from Seurat^42^ was used to calculate the module score for each gene module for 5,000 down-sampled cells which reflects expression of these meta-programs.

### Spatial transcriptomics analysis

Gene/cell count matrices were loaded into Seurat^42^ with spatial information incorporated into the metadata. Transformation and normalisation were performed with SCTransform followed by PCA and UMAP dimensionality reduction. Clustering on the 30 principal components was performed with a resolution of 0.8 to identify clusters of spots. These were visualised spatially with the SpatialDimPlot function. Identification of tumour and normal spots was attempted by genotyping haplotype counts as described above. Deconvolution of Visium spots was performed using the R package CARD using default settings with scRNAseq expression profiles used as a reference^25^. Single cell resolution mapping was then performed with the CARD_SCMapping function with numCell set to 20.

InferCNV (v1.0.2) was used to derive relative copy number difference intensities for each expression-based cluster of spots on each slide^43^. The slide from R1 was used as a reference with the same settings as were used for scRNAseq data. Note the reference R1 spot did not contain any spots with relative copy number changes, confirming this slide indeed displayed uniform copy-number profiles. The scDNAseq subclone with a matching copy number profile for spots from the R1 slide was identified by assessing relative changes which distinguish between R1 subclones but are clonal in other regions. For example, a relative gain in chr10p was not inferred in any other region, therefore this slide does not contain the R1_1 subclone identified in scDNAseq. Relative losses in chr1p were seen in the R2, R3, and R4 slides but not Pa, Pb, R5a, or R5b, suggesting the slide was not from R1_3. Finally, a relative gain in chr5 was seen for R2 and R3 but not Pa or Pb suggesting the R1 slide consisted of a subset of cells from the R1_2 subclone (**Fig. 1e**, **Fig. 4b**, **Supplementary Fig. 19**). The consensus copy number profile was generated by cutting the hierarchical clustering tree of R1_2 subclone generated with the Ward.D2 method into two, selecting the cluster without the loss of part of chr1q (as no relative gains were seen in any other samples) and taking the mode of the copy number state for each genomic bin.

The mean relative inferCNV intensities for each cluster of spots was calculated and centred around zero for each segment, as defined by the shared breakpoints determined from scDNAseq. This intensity was then normalised by the number of copies of R1 for that segment. The normalised expression was then discretized into five different states: gain of one copy, gain of two copies, no event, loss of one copy and loss of two copies. As these slides came from the same tumour and subclones only differ in copy number to a small degree, these five states would cover most of the relative differences from the R1_2 subclone. Inferred total copy number profiles were then calculated by adding the relative discretized gains or losses to the reference R1_2 subclone copy number profile. Total copy number profiles calculated for Visium consensus cluster profiles were then projected onto the scDNAseq UMAP. MEDICC2 was used to derive phylogenetic trees for each slide using the total copy number profiles inferred from Visium clusters^19^. These were then overlaid on the slide as for the LCM data.

## Code availability

Code to generate the main figures is available at GitHub: https://github.com/VanLoo-lab/mpnst_phase_1.

## Data availability

Sequencing data generated in this study have been deposited in the European Genome–phenome Archive (EGA) under accession codes EGAD00001006786 (G&T-seq) and EGAC50000000904 (remaining datasets). Processed data are available at https://doi.org/10.5281/zenodo.19653314.

## Acknowledgements

The authors would like to acknowledge the Genomics Science Technology Platform at the Francis Crick Institute, and in particular Deb Jackson, Marg Crawford, Sam Jones, Hubert Slawinski, and Emma Bourne, for their expertise and support in sequencing the samples used in this study. We would also like to acknowledge the Sanger Single Cell Genomics Core Facility for assistance with G&T-seq data generation, and Mark S. Hill and Emilia L. Lim for assistance with DLP+ analysis pipelines. This work was supported by The Francis Crick Institute, which receives its core funding from Cancer Research UK (CC2008), the UK Medical Research Council (CC2008) and the Wellcome Trust (CC2008). This project was enabled through access to the MRC eMedLab Medical Bioinformatics infrastructure, supported by the Medical Research Council (grant number MR/L016311/1). M.T. was supported as a postdoctoral researcher of the FNRS and a postdoctoral fellow by the European Union’s Horizon 2020 research and innovation program (Marie Skłodowska-Curie Grant agreement no. 747852-SIOMICS). H.Y. was supported by a Crick-CRUK Doctoral Fellowship. J.D. was supported as a postdoctoral fellow of the European Union’s Horizon 2020 research program (Marie Skłodowska-Curie Grant agreement no. 703594-DECODE) and the Research Foundation—Flanders (FWO 12J6916N), and is supported by VIB. A.S. is supported as postdoctoral fellow of the FNRS. R.F.S. is a Professor at the Cancer Research Center Cologne Essen (CCCE) funded by the Ministry of Culture and Science of the State of North Rhine-Westphalia. R.F.S. received support from the German Ministry for Education and Research as BIFOLD - Berlin Institute for the Foundations of Learning and Data (ref. 01IS18025A and ref 01IS18037A). S.Z. was supported by a CRUK Career Development Fellowship (RCCCDF-Nov21\100005). A.M.F. is a NIHR senior investigator and is supported by the National Institute for Health Research, UCLH Biomedical Research Centre and the CRUK Experimental Cancer Centre. P.V.L. is a CPRIT Scholar in Cancer Research and acknowledges CPRIT grant support (RR210006). Figure 1a and 2a in this manuscript were created with BioRender.com, using licenses https://app.biorender.com/illustrations/63e3d78bf737a0fffa159447 (Figure 1a) and https://app.biorender.com/illustrations/63e25d381e4ad47d2694ba37 (Figure 2a).

## Author contributions

A.V., C.C.M., S.W., and T.V. performed tumour profiling experiments, under supervision of P.V.L., A.M.F., S.Z., C.S., and M.T.; H.Y. performed the majority of bioinformatics analyses, with important contributions from Y.C., Y.P., A.S., T.L. and T.K., and with supervision from P.V.L., M.T., S.Z. and R.S.; H.Y. developed the interactive R Shiny app. P.V.L and M.T. designed the study and supervised the project. H.Y., J.D., M.T., and P.V.L. wrote the paper, with important input from Y.C., Y.P., and S.Z. All authors read and approved the final version of the manuscript.

## Conflicts of interests

C.S. acknowledges grant support from Pfizer, AstraZeneca, Bristol Myers Squibb, Roche-Ventana, Boehringer-Ingelheim, Archer Dx Inc (collaboration in minimal residual disease sequencing technologies) and Ono Pharmaceutical, is an AstraZeneca Advisory Board member and Chief Investigator for the MeRmaiD1 clinical trial, has consulted for Pfizer, Novartis, GlaxoSmithKline, MSD, Bristol Myers Squibb, Celgene, AstraZeneca, Illumina, Genentech, Roche-Ventana, GRAIL, Medicxi, Bicycle Therapeutics, and the Sarah Cannon Research Institute, has stock options in Apogen Biotechnologies, Epic Bioscience, GRAIL, and is co-founder and has shares of Achilles Therapeutics. All other authors declare no conflicts of interest.

## Supplementary figures

**Supplementary Figure 1.**
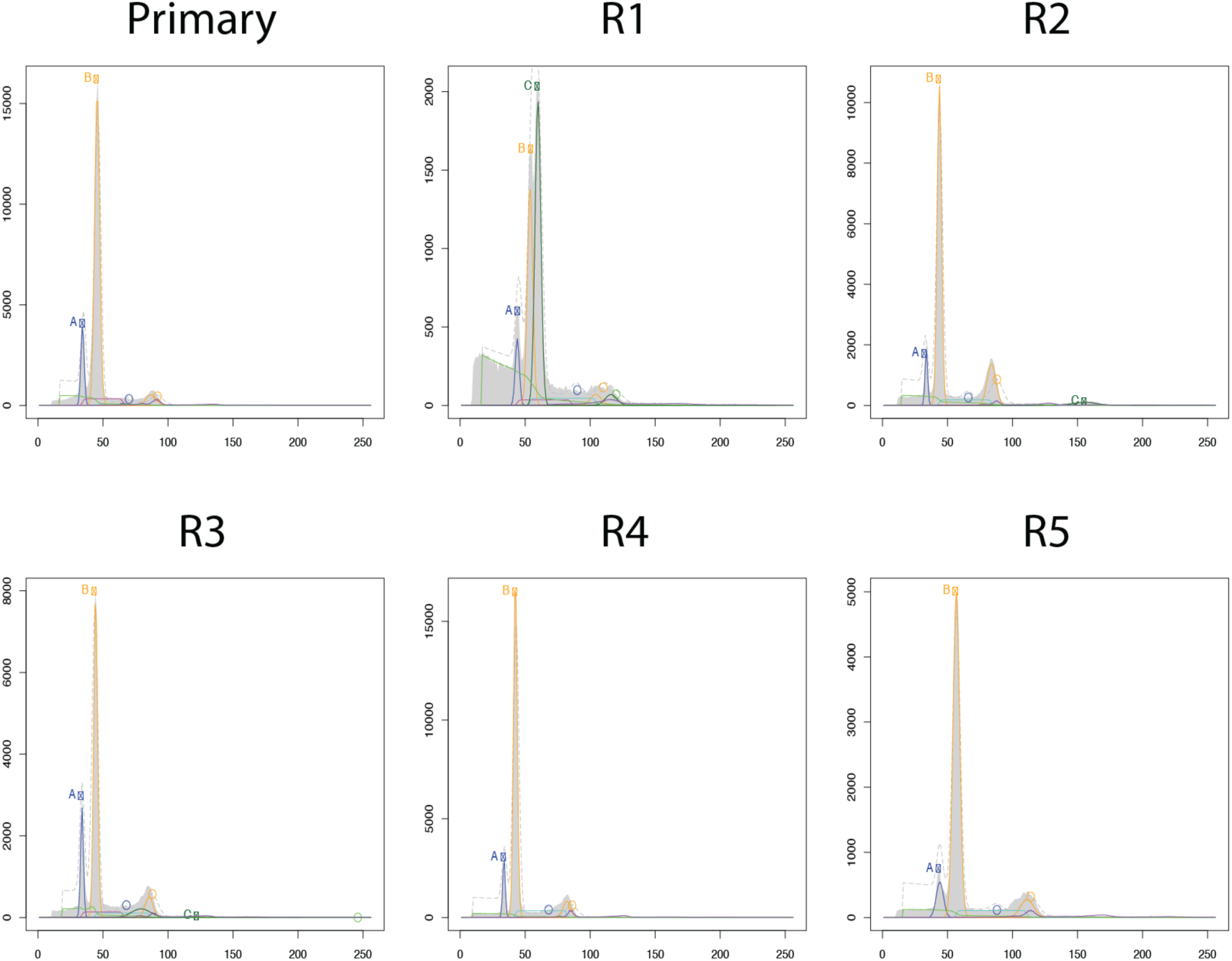
FACS ploidy analysis of primary and recurrent tumours. Histograms (PI-Area vs Count) show DNA content distributions of nuclei isolated from the primary and five recurrent lesions (R1–R5), analysed by flow cytometry. Nuclei were gated by size and granularity (forward scatter, FSC *vs.* side scatter, SSC), and single nuclei were selected based on area *vs.* height parameters. Blue peaks (A): Diploid (2N) population; Yellow peaks (B) and Dark Green peaks (C): Aneuploid populations. Light green: cell debris.

**Supplementary Figure 2.**
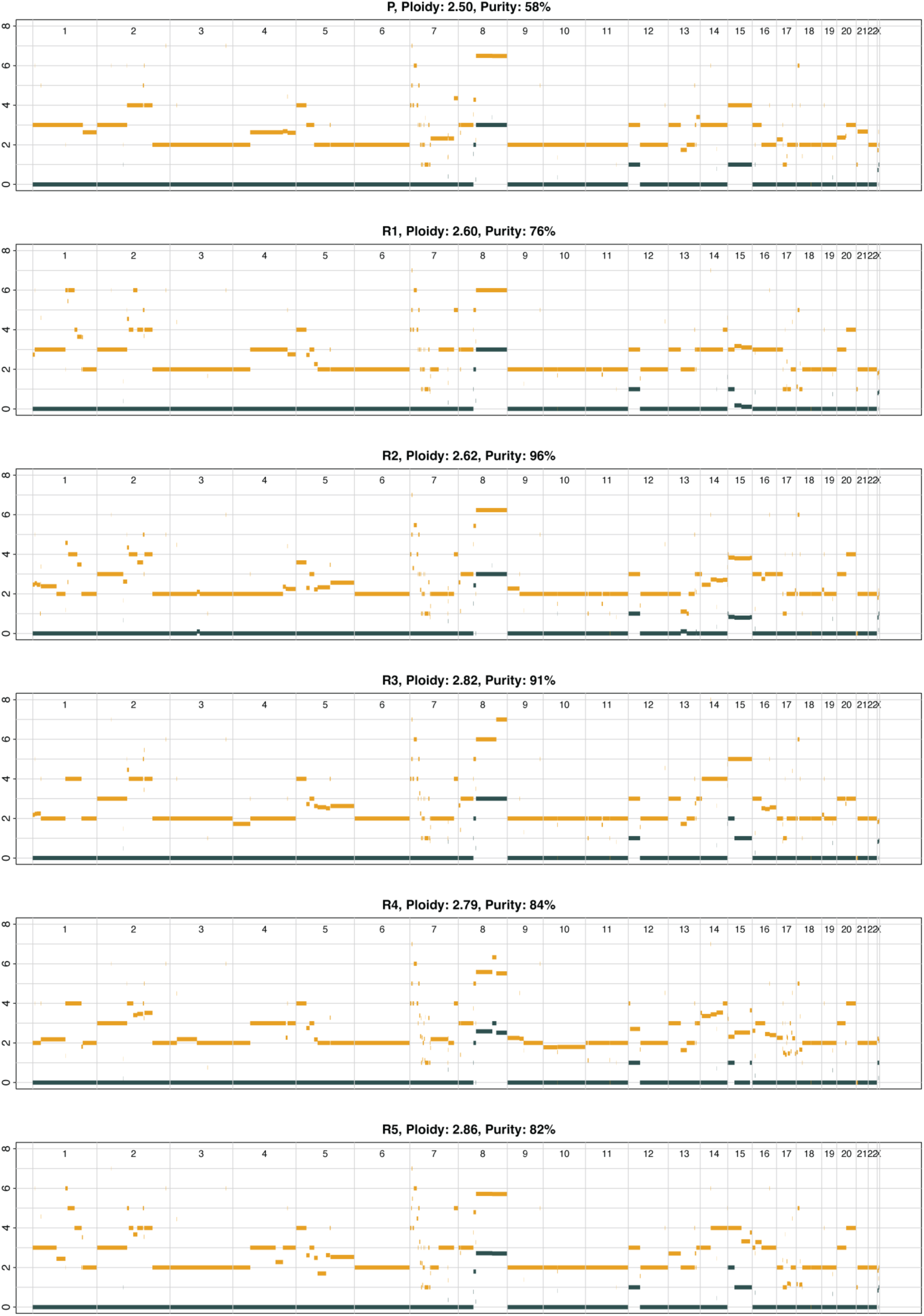
Battenberg copy number profiles per region. Allele-specific copy number for all chromosomes with total copy number shown in orange and the minor alleles in grey, supporting Fig. 1b. Non-integer values indicate the presence of multiple subclones with different copy number states.

**Supplementary Figure 3.**
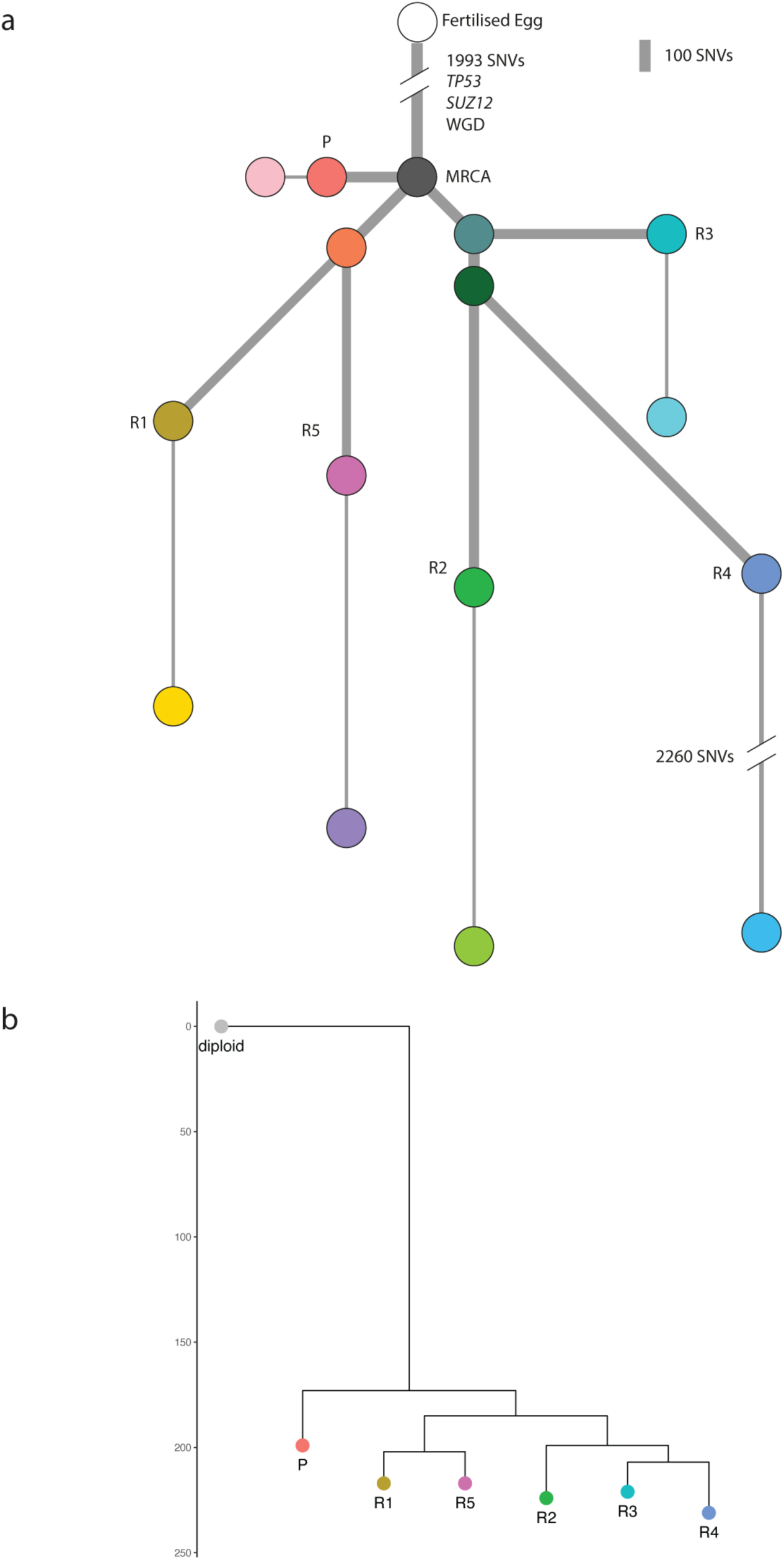
SNV and CNA-based phylogenetic trees. (**a**) Reconstructed SNV-based clone tree derived from bulk WGS data using ndDPClust. The length of the branches is proportional to number of SNVs. Truncal homozygous deletion of *SUZ12*, frameshift deletion of *TP53*, and WGD status were annotated. (**b**) Reconstructed CNA-based sample tree derived using MEDICC2 analysis of bulk copy number profiles. The length of branches indicates the number of copy number events.

**Supplementary Figure 4.**
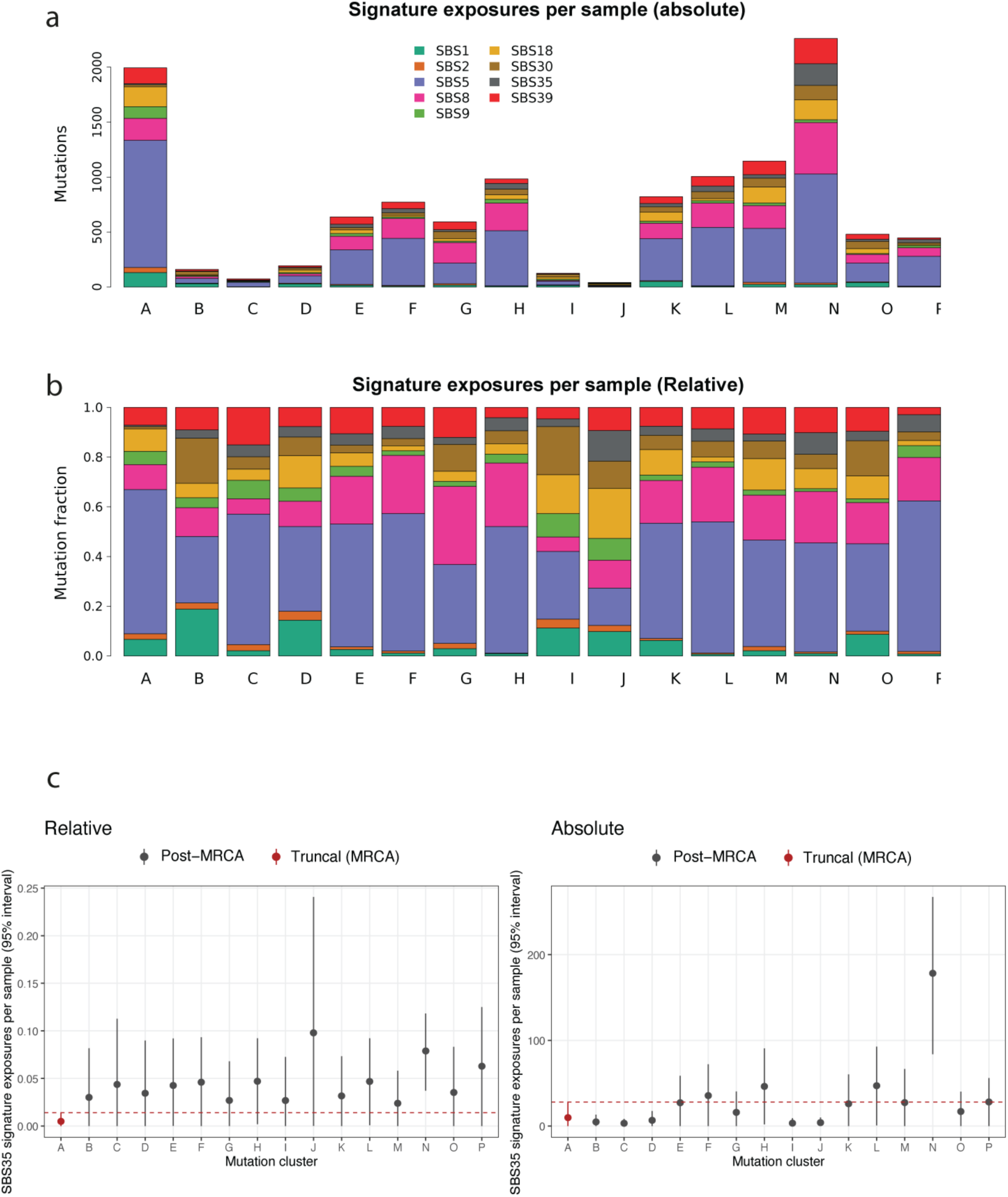
Mutational signature profiles among bulk tree branches. Absolute (**a**) and relative (**b**) exposures of COSMIC mutational signatures in each mutational cluster letter-coded as in Fig. 1c. (c) 95% confidence interval of SBS35 signature exposures per sample.

**Supplementary Figure 5.**
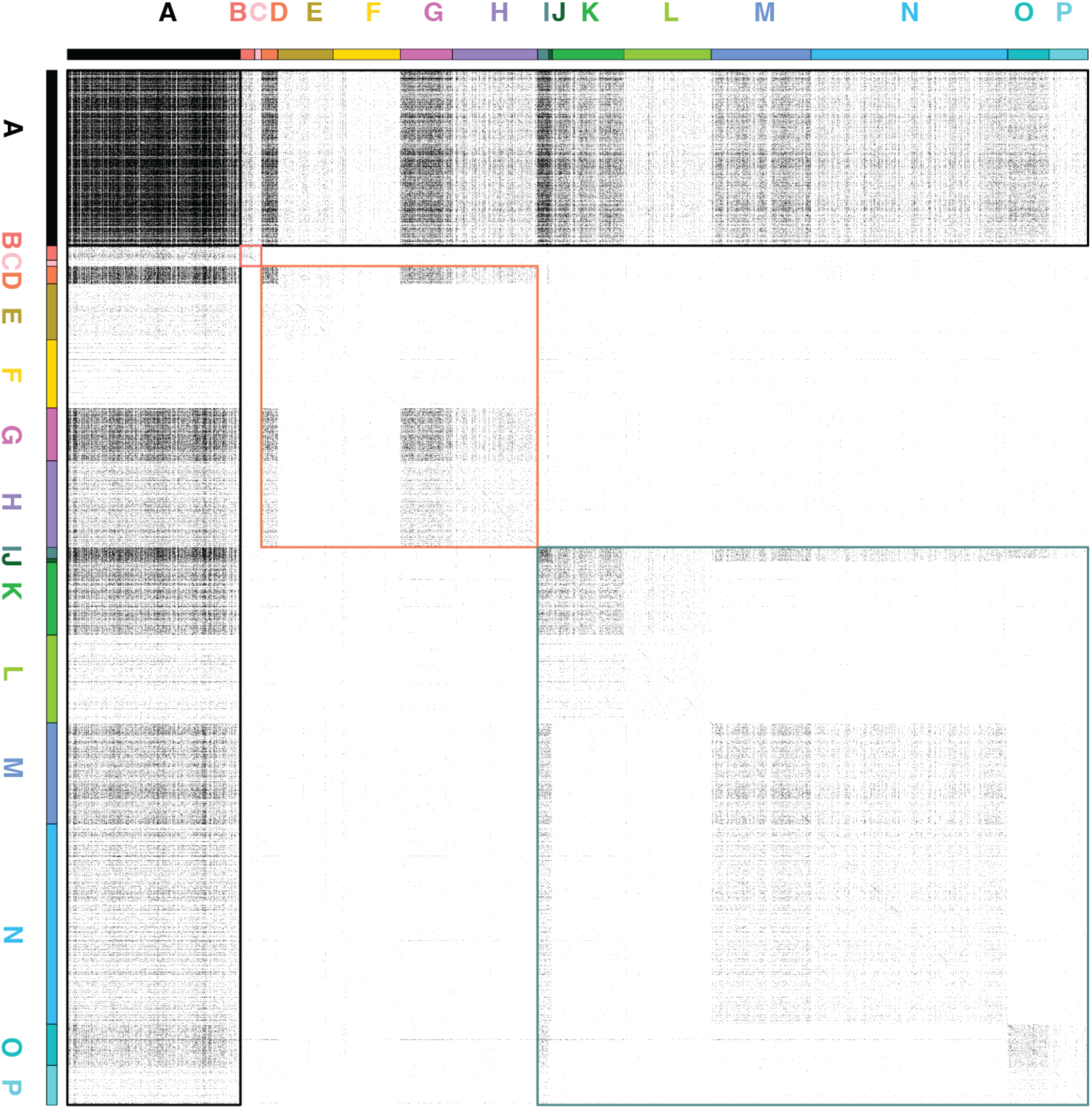
Co-occurrence matrix of bulk-detected SNVs in scDNAseq data. Mutations are aggregated in mutation clusters identified by ndDPClust. Colour-code of clusters along the axes follows Fig. 1c. Different colour squares indicate prominent co-occurrence patterns following major clades in the phylogenetic tree: clonal mutations co-occur with mutations in all clusters (black); mutations specific to the primary tumour only co-occur with other mutations specific to the primary tumour (red); mutations specific to the R1-R5 clade only co-occur with other mutations specific to the R1-R5 clade (orange), and follow further phylogenetic relationships within that clade; mutations specific to the R2-R3-R4 clade only co-occur with other mutations specific to the R2-R3-R4 clade (green), and follow further phylogenetic relationships within that clade.

**Supplementary Figure 6.**
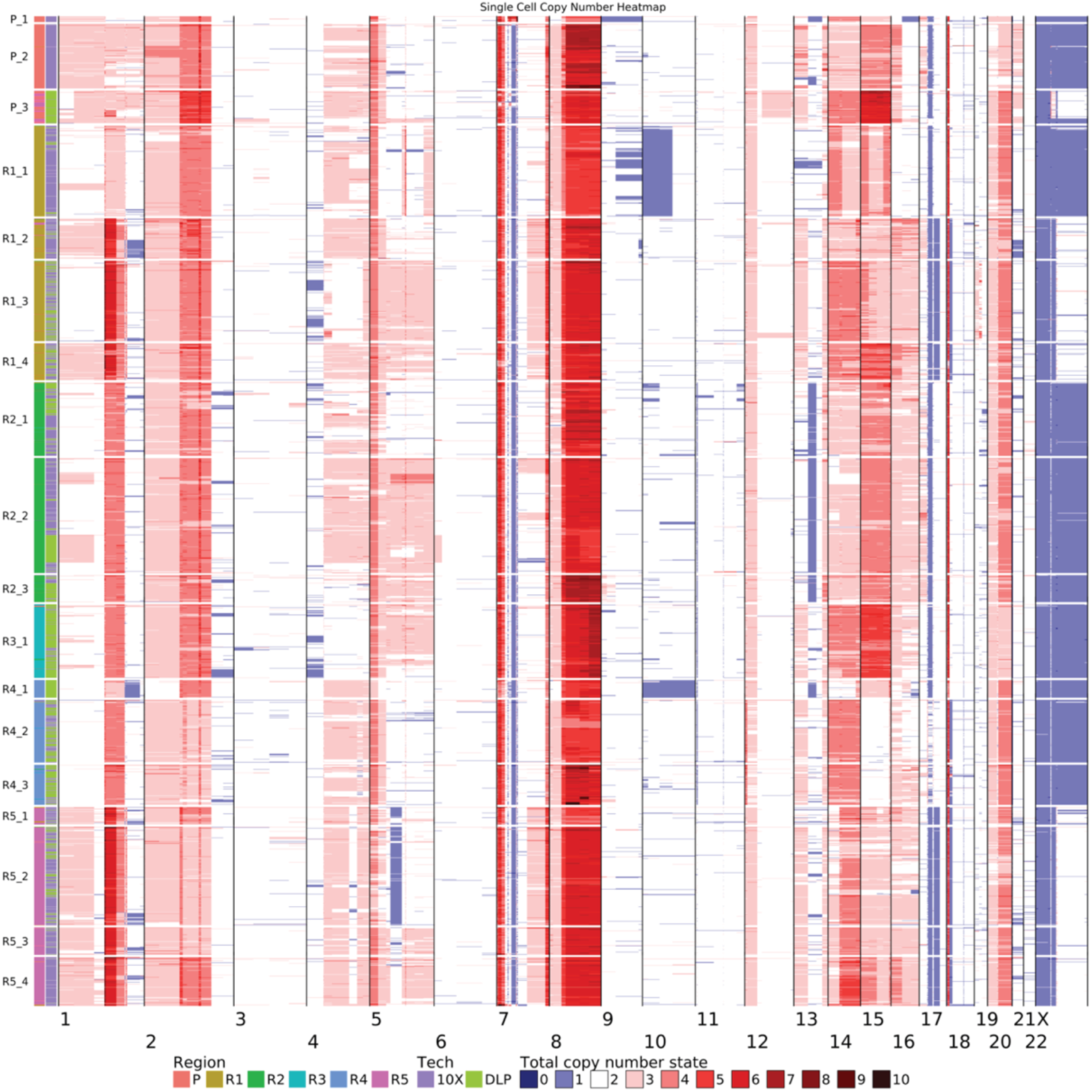
Total copy number profiles of scDNAseq tumour cells. Heatmap showing the total copy number profiles of scDNAseq tumour cells. 18 subclones were defined using K-means clustering approach.

**Supplementary Figure 7.**
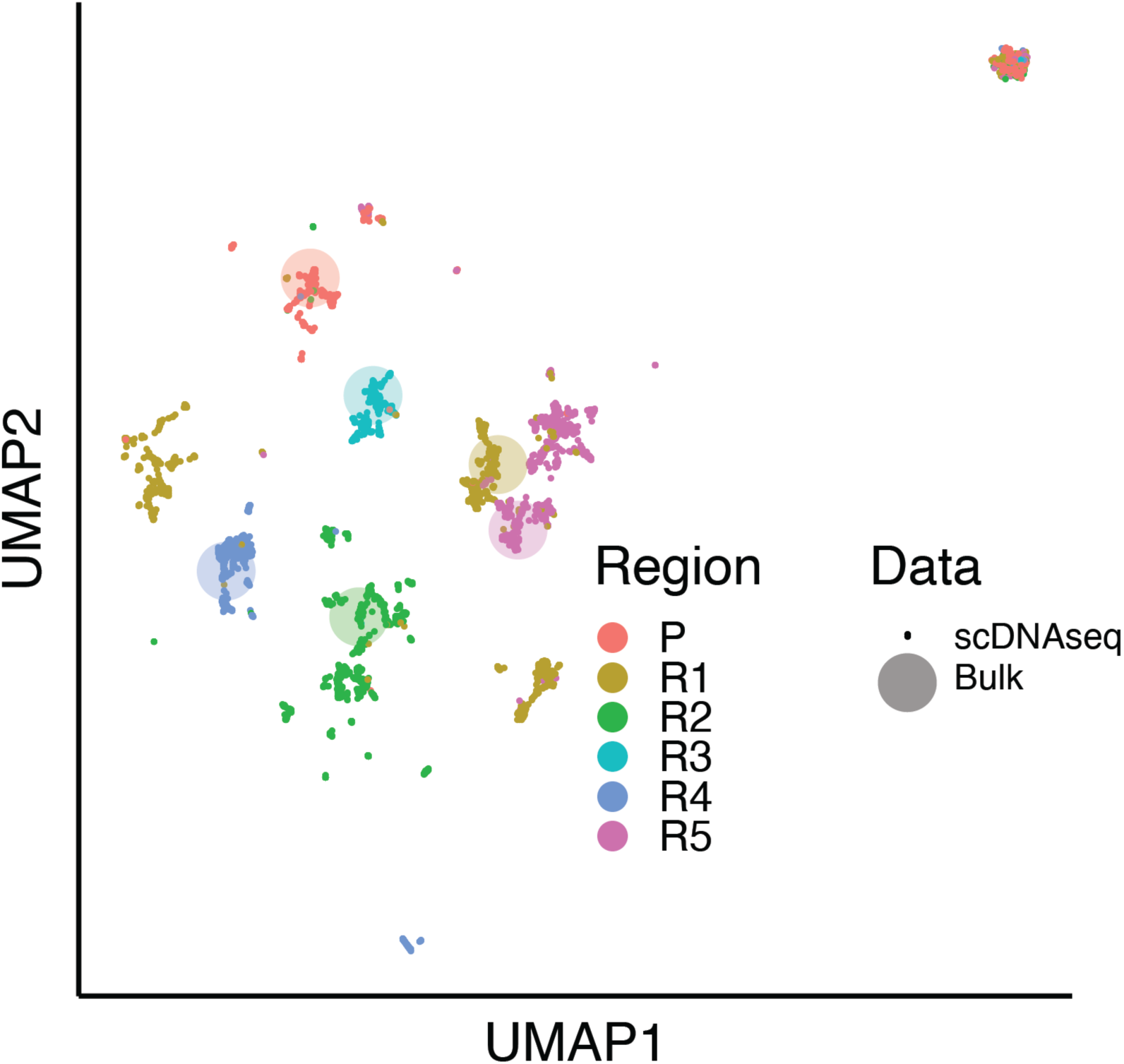
UMAP projection of CNA profiles from scDNAseq cells and bulk WGS. Cells from scDNAseq are shown as points and bulk profiles are shown as larger circles. Both are annotated by their region of origin.

**Supplementary Figure 8.**
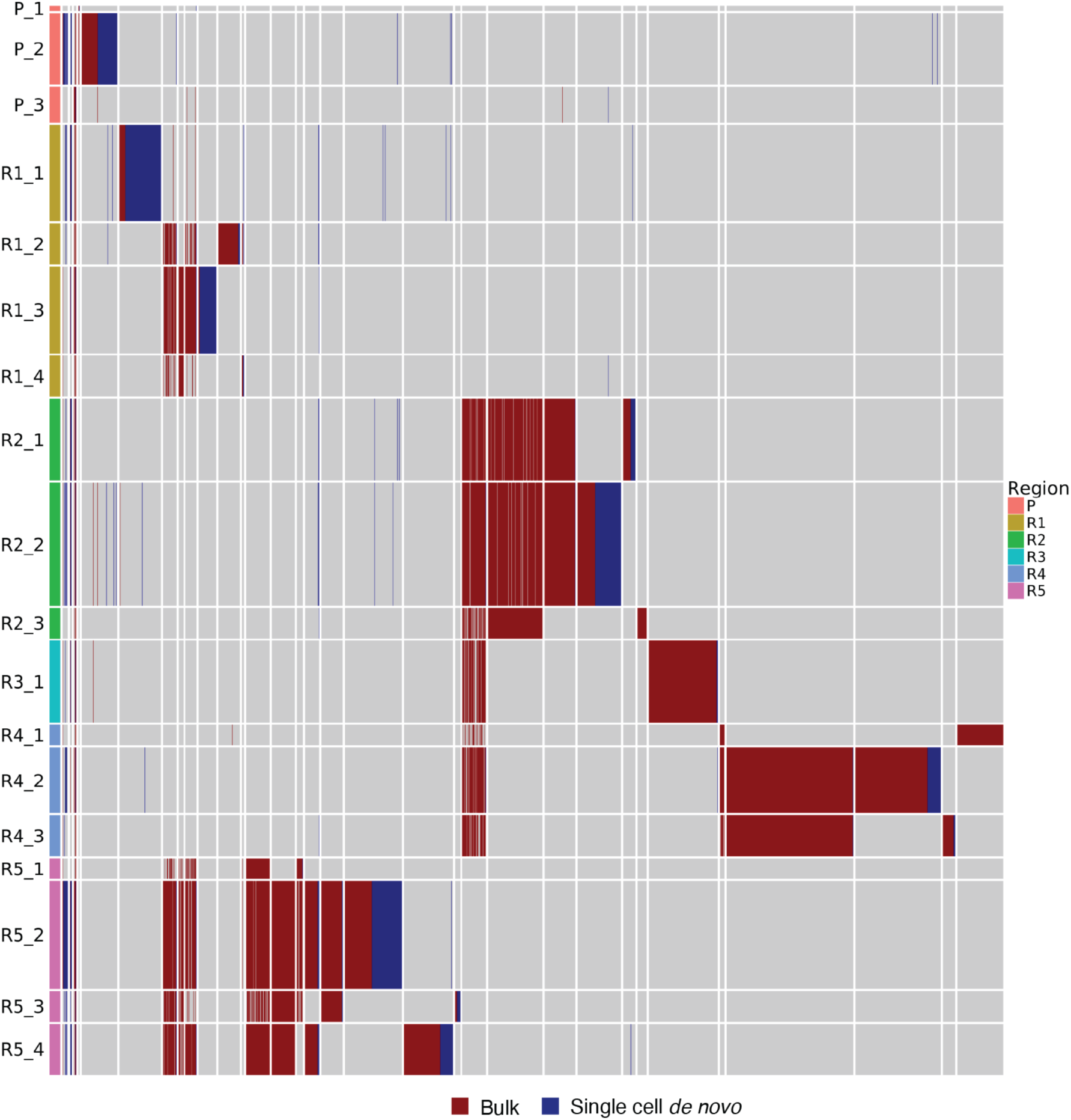
Results from genotyping of SNVs called from bulk WGS and pseudobulk derived from scDNAseq. SNVs called in the bulk and *de novo* calls from scDNAseq are shown in red and blue, respectively. Row height is proportional to the total read depth of each cluster of cells.

**Supplementary Figure 9.**
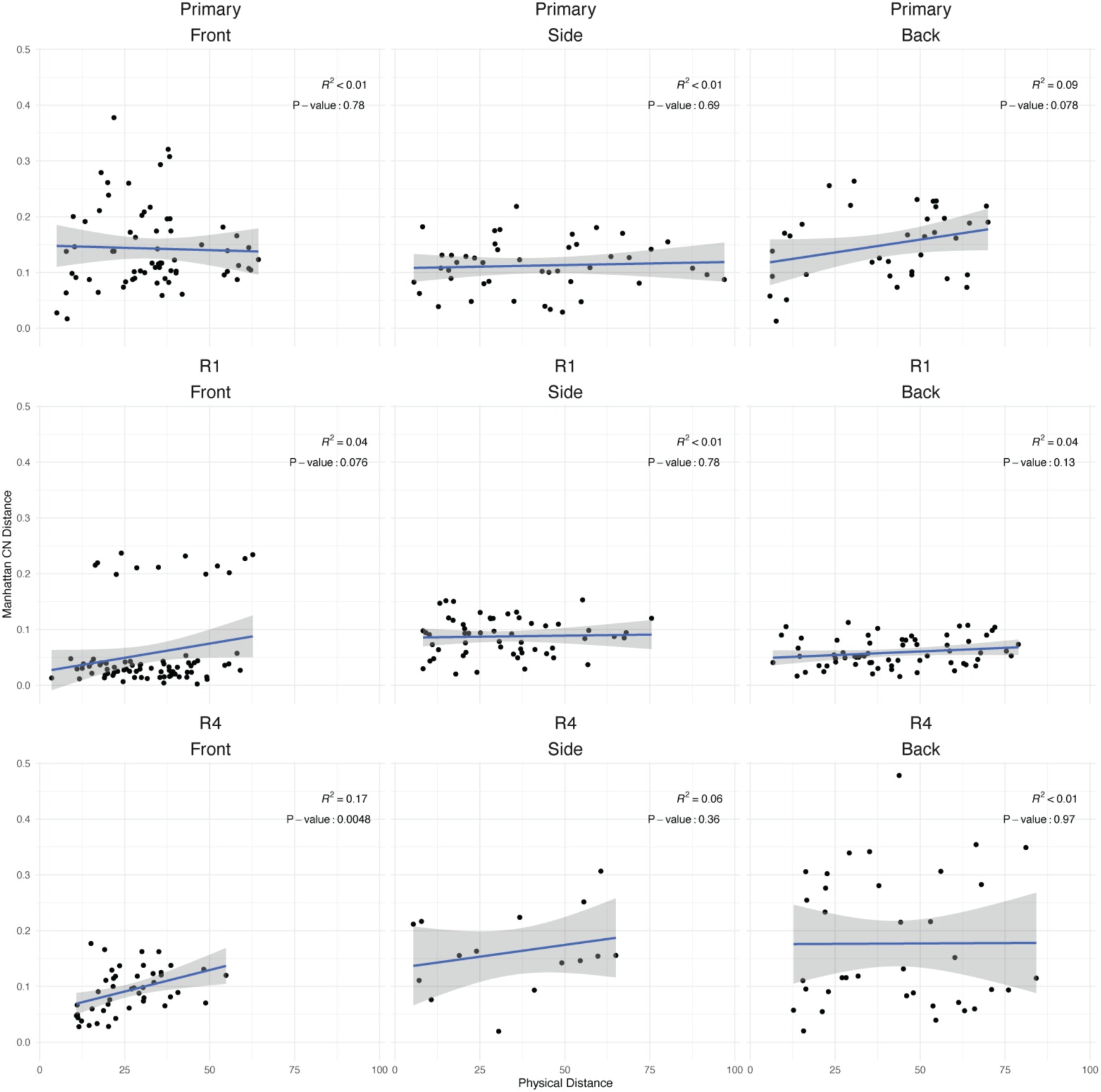
Correlation of physical and genetic distances in LCM sections. Allele-specific CNA profiles were derived from mini-bulk LCM samples. The correlation between physical distances (Euclidean distances from images) and genetic distances (Manhattan distances of copy number profiles) between spots are shown.

**Supplementary Figure 10.**
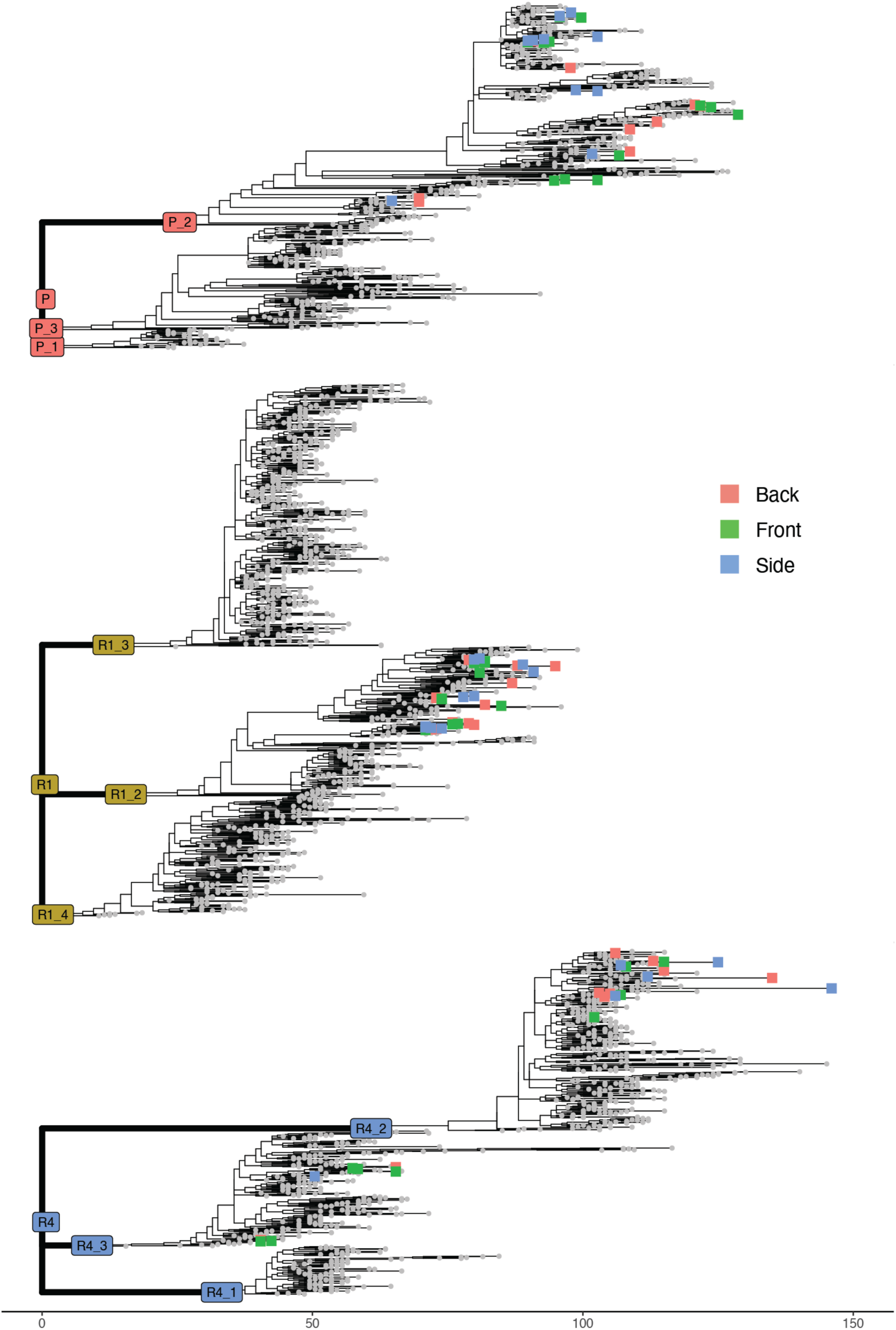
Phylogenetic analysis of LCM mini-bulk data. LCM spots highlighted on phylogenetic trees shown in Fig. 2c, colour-coded by section origins (front, side, or back).

**Supplementary Figure 11.**
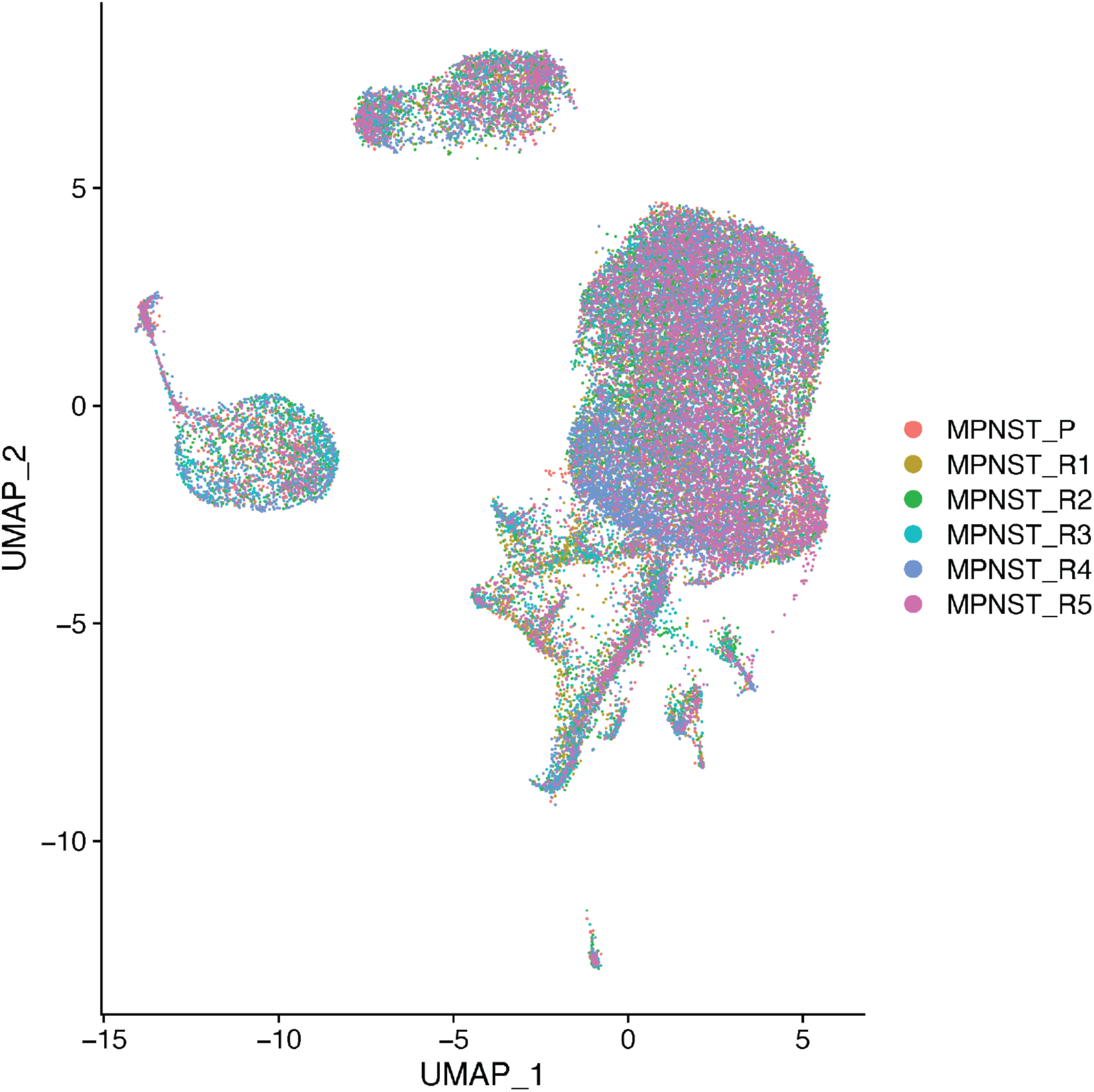
Batch correction removes transcriptomic effects of CNAs. UMAP projection after batch effect correction, annotated by sample origin. Cells from all tumour regions overlap and the biological effects of CNAs on gene expression are overcorrected as technical noise (**Methods**).

**Supplementary Figure 12.**
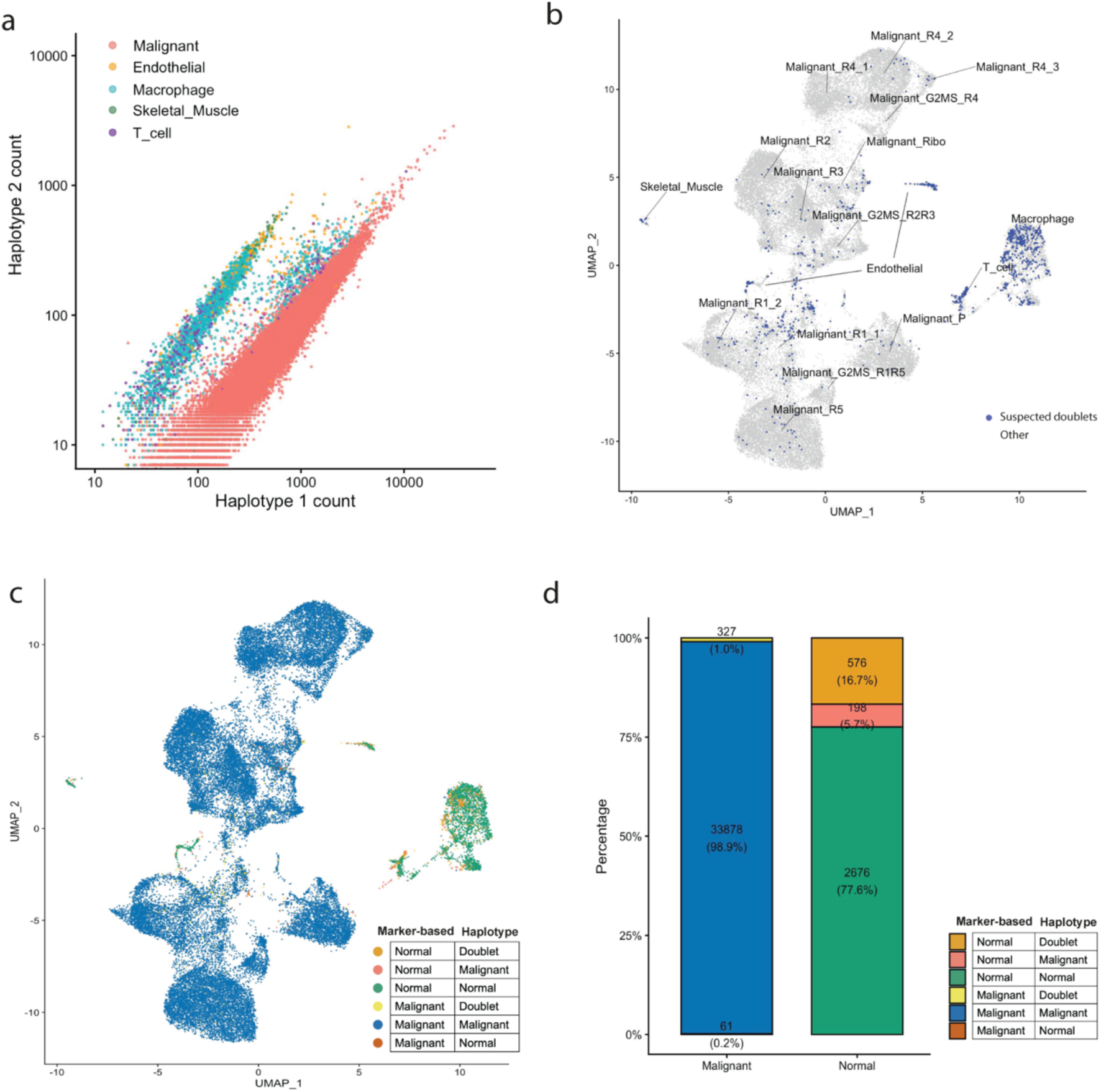
Haplotype-based orthogonal validation of scRNAseq cell-type annotation and doublet detection. (**a**) Cells from Fig. 3b, re-coloured by cell type annotation. (**b**) A subset of scRNAseq cells identified as putative doublets by haplotype are highlighted (navy) on the scRNAseq UMAP projection, with simulated doublets used as a reference. (**c**) UMAP projection colour-coded by haplotype-based (Normal/Tumour/Doublet) annotation, quantified as the confusion matrix in (**d**).

**Supplementary Figure 13.**
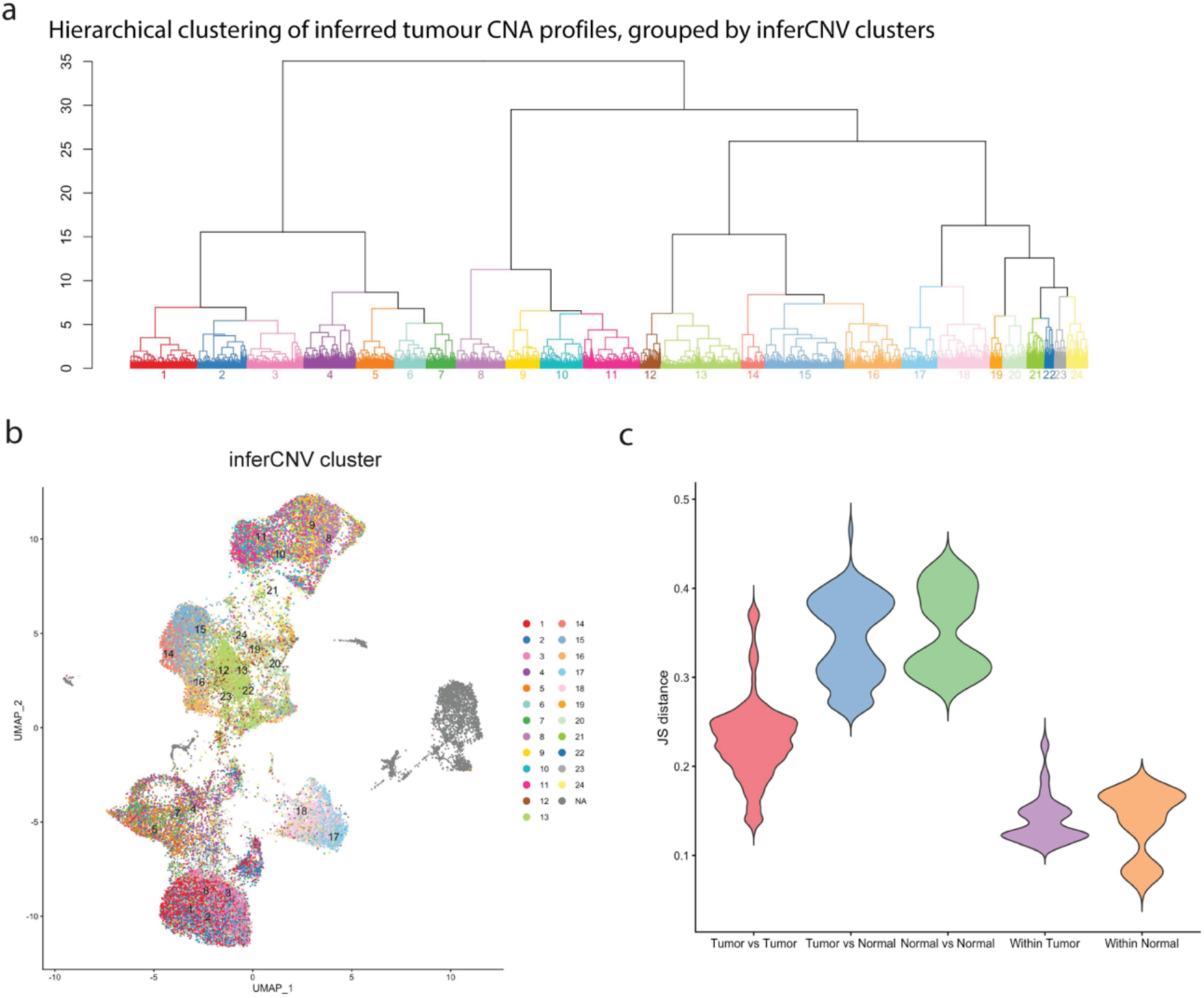
CNA-driven transcriptomic similarity between scRNAseq cells assessed by inferCNV clustering and Jensen-Shannon distances. (**a**) Hierarchical clustering of tumour CNA profiles derived from inferCNV. (**b**) UMAP projection of scRNAseq data, colour-coded by inferCNV clusters. (**c**) Pairwise Jensen-Shannon distances between pseudobulk transcriptomic profiles of tumour clusters defined by inferCNV and normal cell types annotated by canonical markers. Within-group comparisons serve as baselines.

**Supplementary Figure 14.**
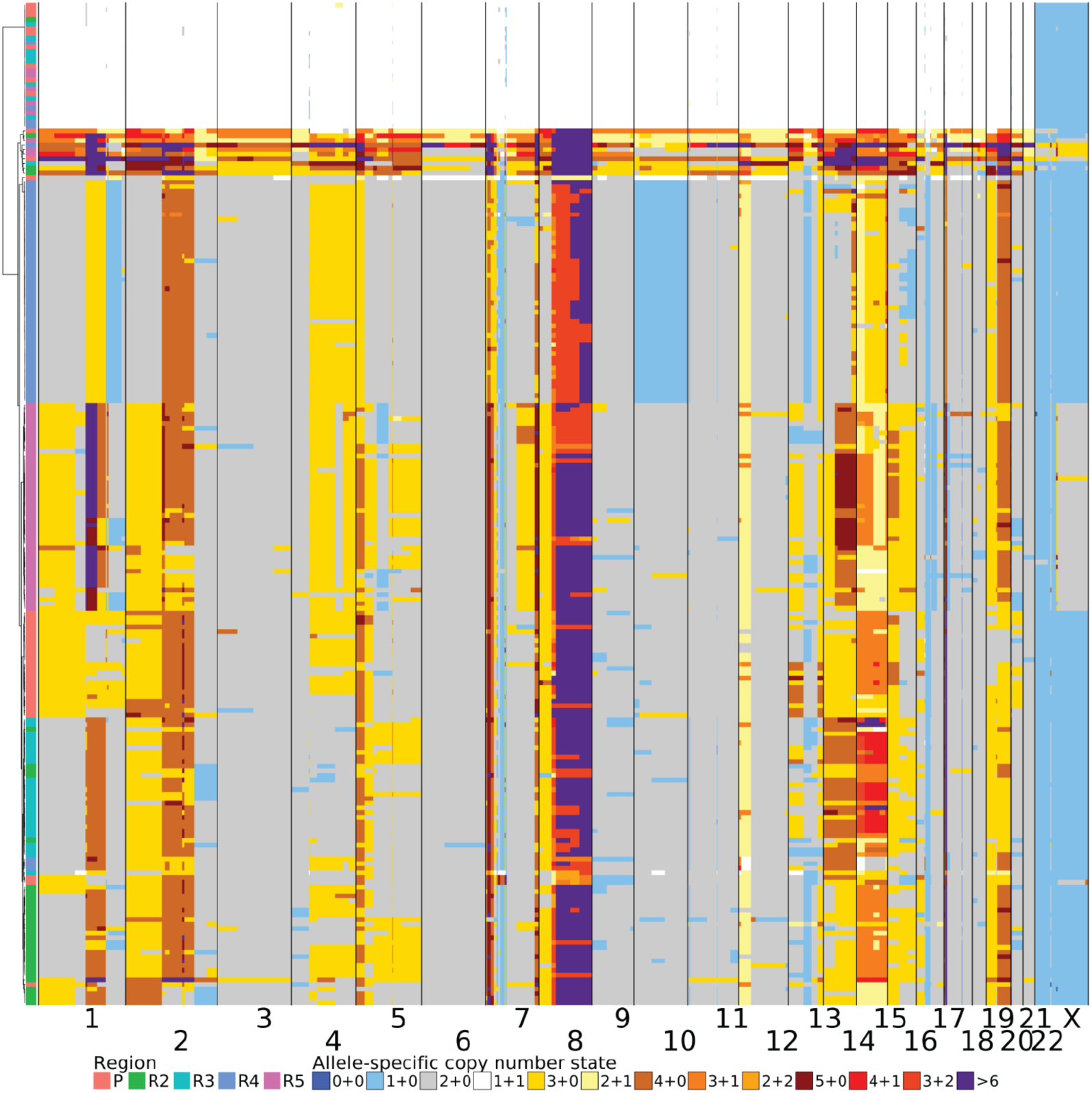
Allele specific copy number profiles derived from the DNA layer of G&T-sequenced cells. Allele-specific copy number heatmap derived from 216 single-cell genomes across five samples in G&T-seq.

**Supplementary Figure 15.**
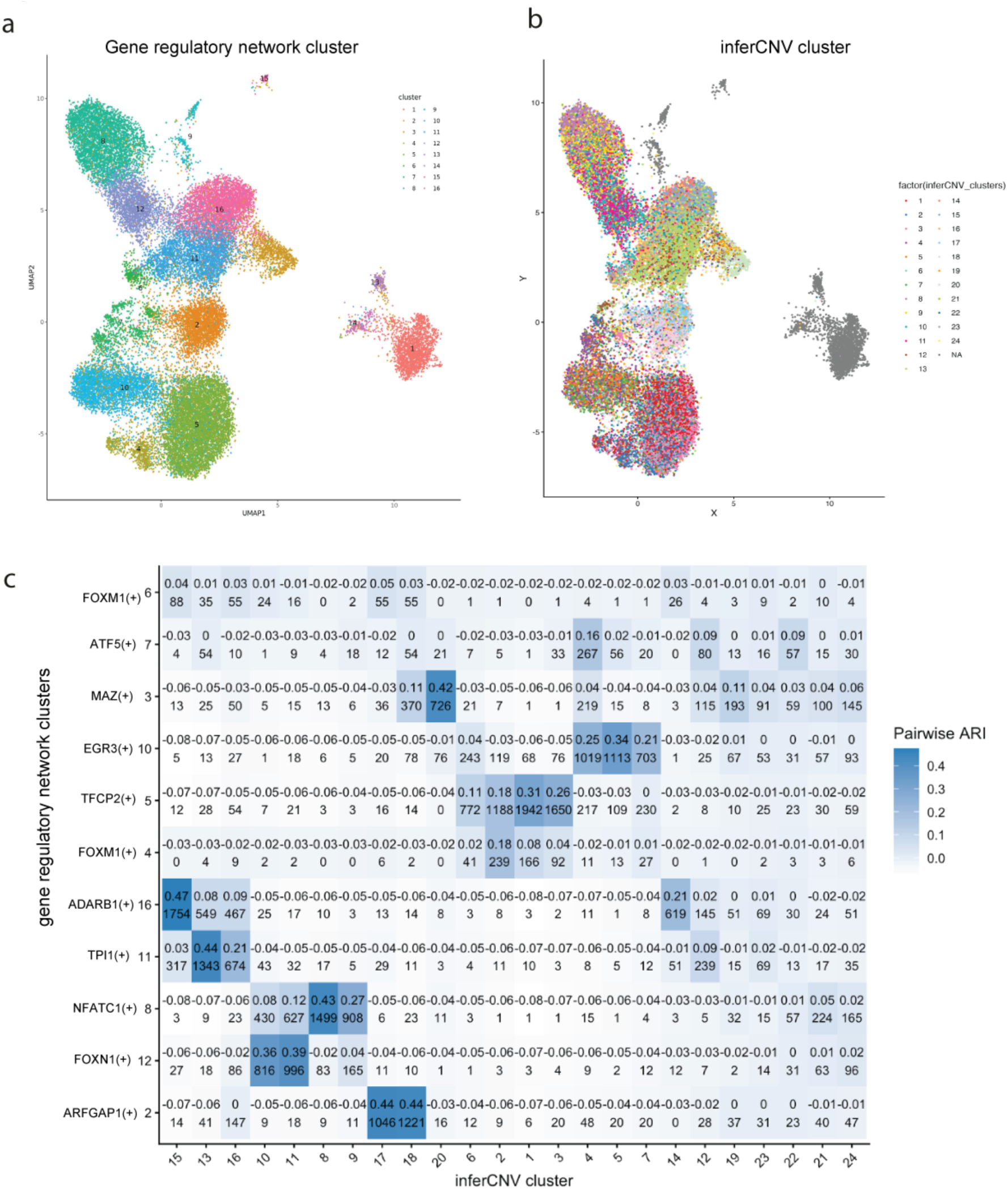
Gene regulatory network UMAP projection of scRNAseq cells. (**a**) UMAP projection of regulon activity in scRNA-seq cells derived from SCENIC, colour-coded by SCENIC gene regulatory network clusters and (**b**) inferCNV clusters. Normal cells are coloured in grey. (**c**) Pairwise adjusted rand index (ARI) between SCENIC clusters and inferCNV clusters. Cell count for each intersection is shown below the ARI value.

**Supplementary Figure 16.**
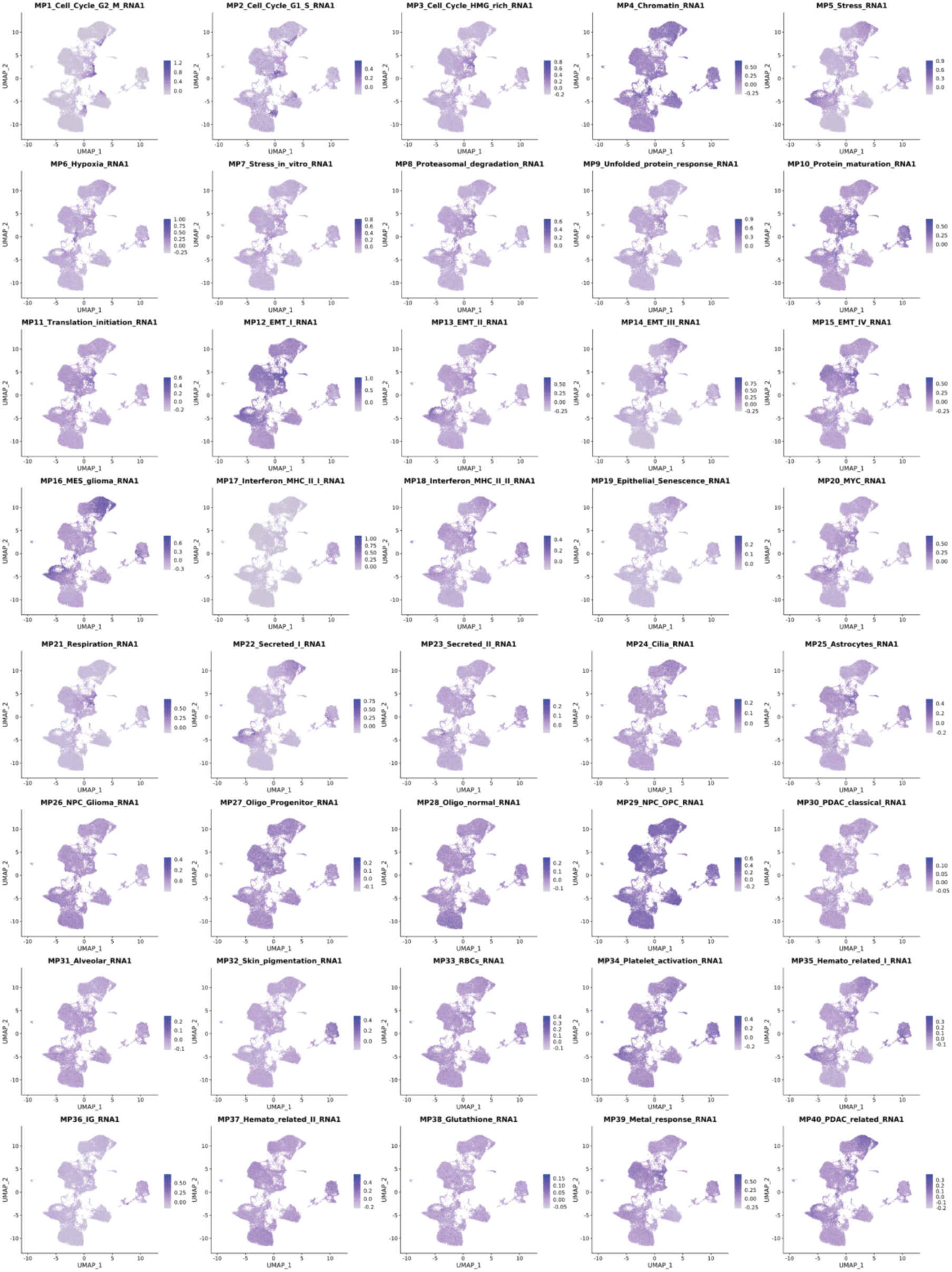
Activity scores for cancer cell states. UMAP visualisations showing the distribution of module scores for pan-cancer meta-programs. Relative expression levels are shown with a light grey to dark blue gradient for low to high activity.

**Supplementary Figure 17.**
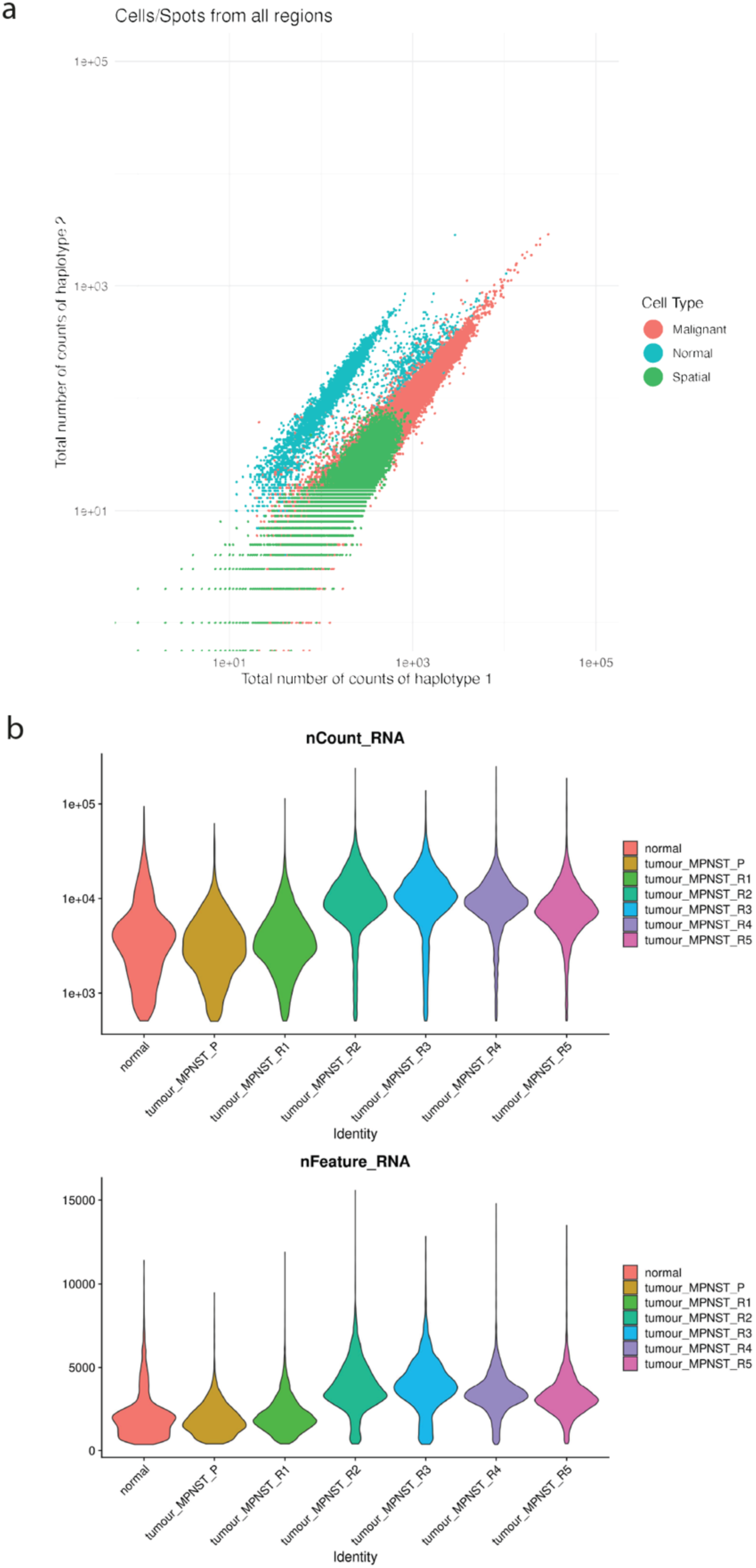
Genotyping and transcript outputs of Visium spots. (**a**) Haplotype counts of spots from spatial transcriptomic slides mapped to Fig. 3b. Haplotype counts of cells from all regions scRNAseq data (blue or red) or spots from all eight Visium slides (green) are shown. (**b**) Violin plots of nCount_RNA (log10-transformed) and nFeature_RNA across cell types in scRNAseq data, showing higher number of transcripts captured in malignant cells from recurrence regions compared to normal cells.

**Supplementary Figure 18.**
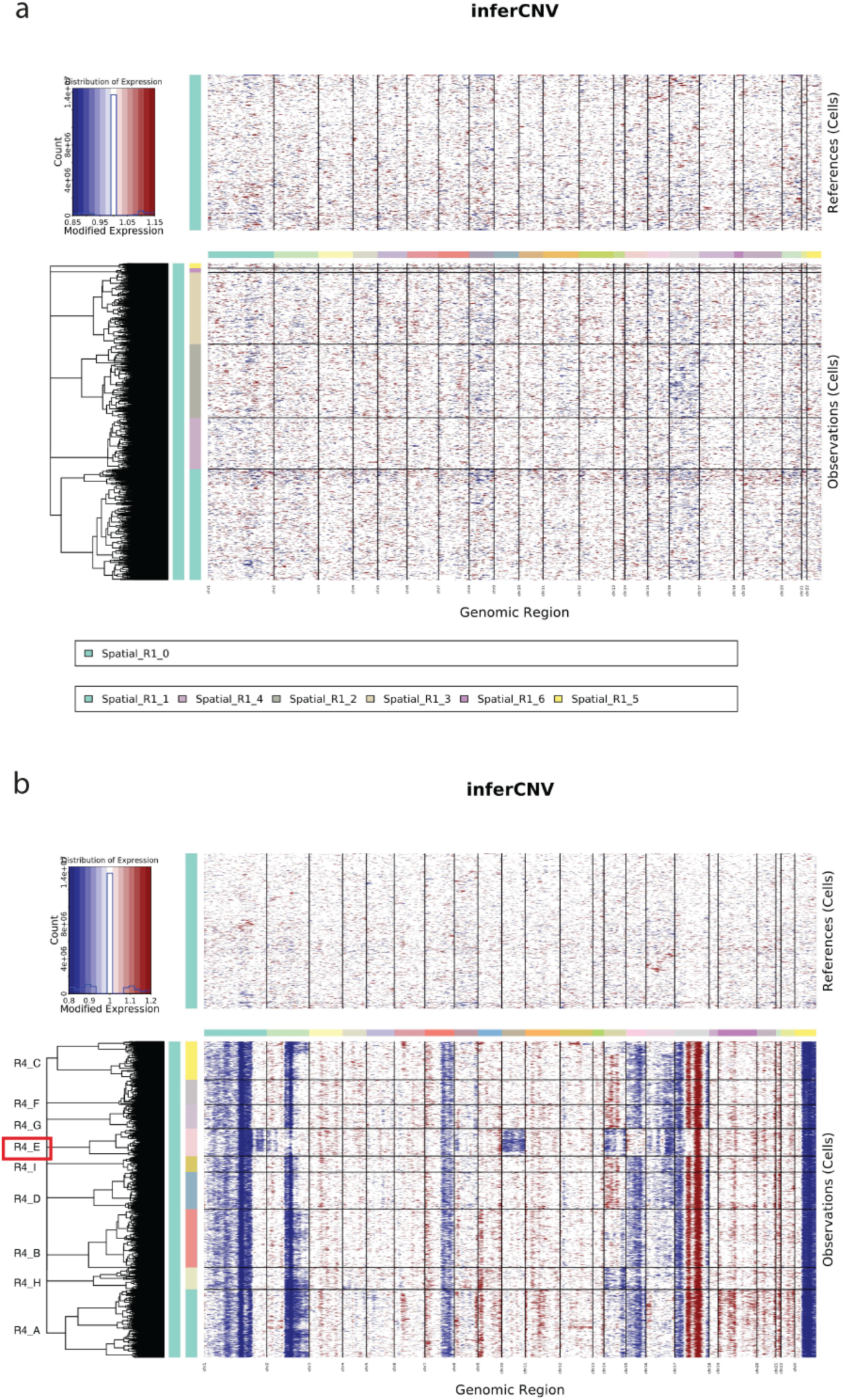
InferCNV profiles for spatial transcriptomic slides using R1 as reference. (**a-b**) InferCNV copy number profiles of (a) R1 and (b) R4, with R1 as reference, which did not display any copy number heterogeneity within the slide. R4 subclones R4_A to R4_I are labelled in (b), with R4_E highlighted, which is a subclone showing copy number loss in chromosome 10.

**Supplementary Figure 19.**
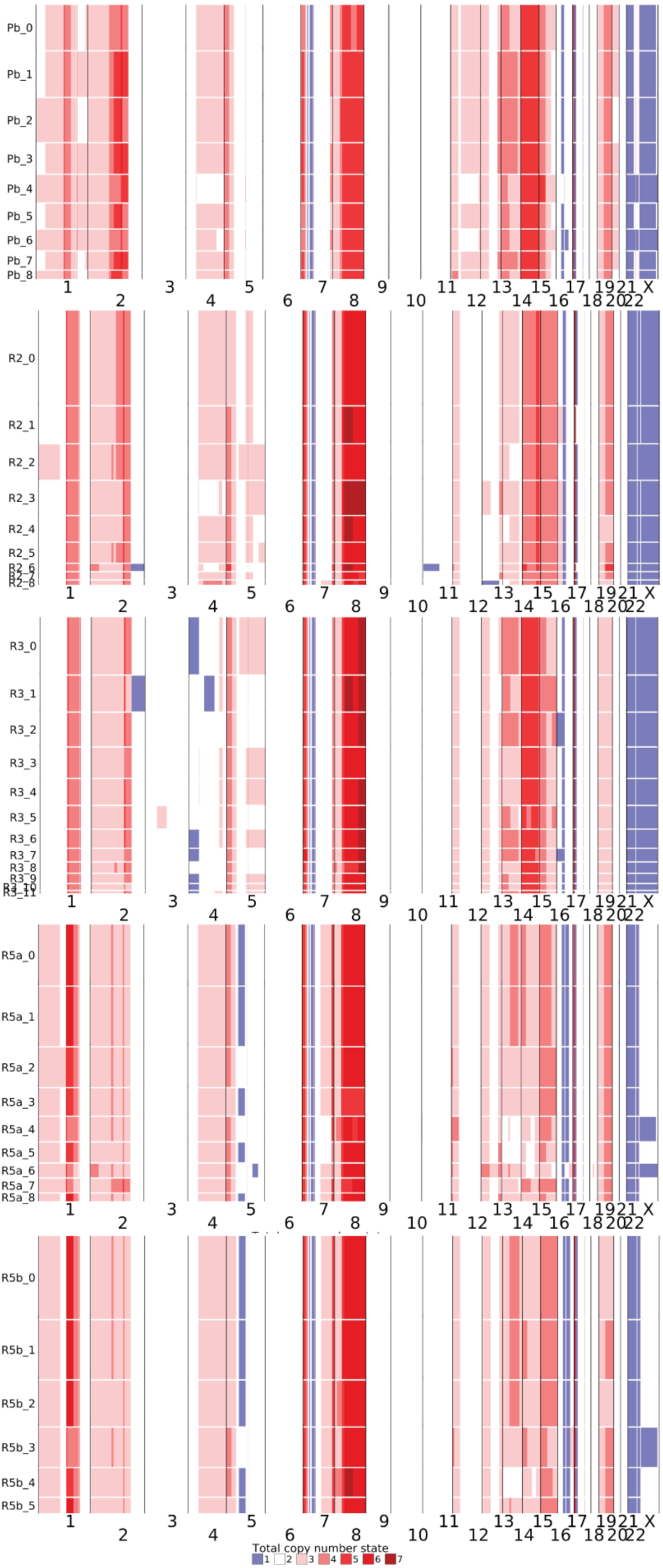
Inferred integer copy-number profiles for Pb, R2, R3, R5a and R5b. Inferred integer copy-number profiles of clusters for Pb, R2, R3, R5a and R5b. R1 was used as reference.

**Supplementary Figure 20.**
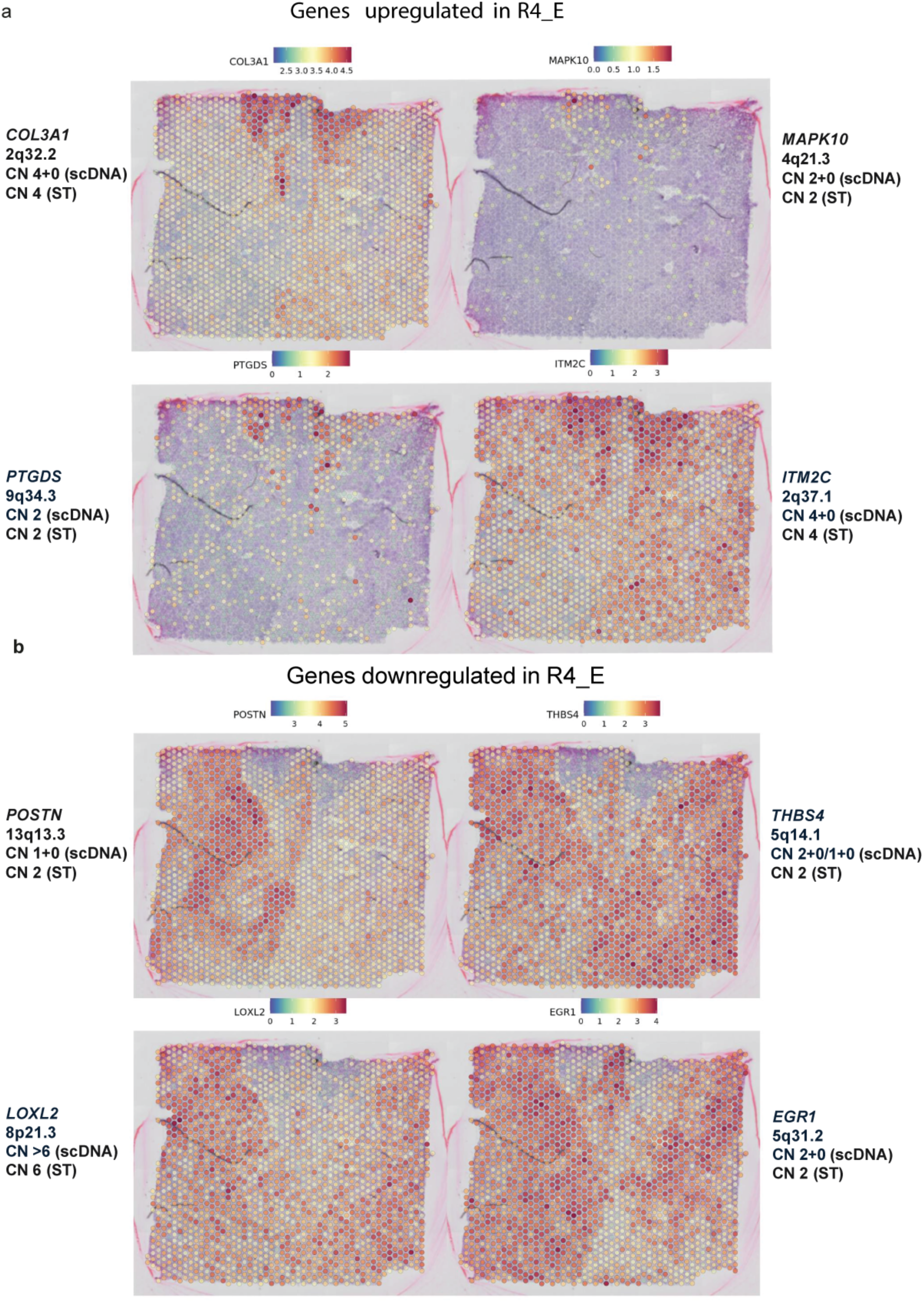
Differentially expressed genes in R4_E showed no specific CNA trend comparing to other R4 regions. (**a-b**) Top genes that were significantly upregulated (a) or downregulated (b) in differential gene expression analysis comparing R4_E and other spots in spatial transcriptomic data. The genome location and copy number status of the genes in scDNAseq and Spatial Transcriptomics are labelled on the plot.

## Supplementary Tables

**Supplementary Table 1.** Cancer cell fraction (CCF) and number of SNVs of each identified cluster across samples.

| Cluster | Primary | R1 | R2 | R3 | R4 | R5 | Number<br>of SNVs |
| --- | --- | --- | --- | --- | --- | --- | --- |
| A | 1.04 | 1.02 | 1.08 | 1.05 | 1.12 | 1.01 | 1993 |
| B | 0.83 | 0.00 | 0.00 | 0.00 | 0.02 | 0.01 | 161 |
| C | 0.26 | 0.00 | 0.00 | 0.00 | 0.00 | 0.00 | 74 |
| D | 0.00 | 0.87 | 0.00 | 0.00 | 0.00 | 0.86 | 194 |
| E | 0.00 | 0.80 | 0.00 | 0.00 | 0.00 | 0.00 | 638 |
| F | 0.00 | 0.25 | 0.00 | 0.00 | 0.00 | 0.00 | 772 |
| G | 0.00 | 0.00 | 0.00 | 0.00 | 0.00 | 0.77 | 593 |
| H | 0.00 | 0.00 | 0.00 | 0.00 | 0.00 | 0.25 | 983 |
| I | 0.00 | 0.00 | 0.95 | 0.94 | 1.01 | 0.00 | 125 |
| J | 0.00 | 0.00 | 1.10 | 0.00 | 1.13 | 0.00 | 41 |
| K | 0.00 | 0.00 | 0.90 | 0.00 | 0.00 | 0.00 | 822 |
| L | 0.00 | 0.00 | 0.28 | 0.00 | 0.00 | 0.00 | 1005 |
| M | 0.00 | 0.00 | 0.00 | 0.89 | 0.00 | 0.00 | 1145 |
| N | 0.00 | 0.00 | 0.00 | 0.38 | 0.00 | 0.00 | 2260 |
| O | 0.00 | 0.01 | 0.00 | 0.90 | 0.00 | 0.00 | 481 |
| P | 0.00 | 0.00 | 0.00 | 0.29 | 0.00 | 0.00 | 447 |

## References

1. Dentro, S.C. et al. Characterizing genetic intra-tumor heterogeneity across 2,658 human cancer genomes. Cell 184, 2239–2254 (2021).

2. Jamal-Hanjani, M. et al. Tracking the Evolution of Non-Small-Cell Lung Cancer. N Engl J Med 376, 2109–2121 (2017).

3. Turajlic, S. et al. Deterministic Evolutionary Trajectories Influence Primary Tumor Growth: TRACERx Renal. Cell 173, 595–610 (2018).

4. Litchfield, K. et al. Representative Sequencing: Unbiased Sampling of Solid Tumor Tissue. Cell Rep 31, 107550 (2020).

5. Minussi, D.C. et al. Breast tumours maintain a reservoir of subclonal diversity during expansion. Nature 592, 302–308 (2021).

6. Barkley, D. et al. Cancer cell states recur across tumor types and form specific interactions with the tumor microenvironment. Nat Genet 54, 1192–1201 (2022).

7. Lucas, O. et al. Characterizing the evolutionary dynamics of cancer proliferation in single-cell clones with SPRINTER. Nat Genet 57, 103–114 (2025).

8. Lomakin, A. et al. Spatial genomics maps the structure, nature and evolution of cancer clones. Nature 611, 594–602 (2022).

9. Erickson, A. et al. Spatially resolved clonal copy number alterations in benign and malignant tissue. Nature 608, 360–367 (2022).

10. Sun, D. et al. Stem-like cells drive NF1-associated MPNST functional heterogeneity and tumor progression. Cell Stem Cell 28, 1397–1410 (2021).

11. Dunn, G.P. et al. Role of resection of malignant peripheral nerve sheath tumors in patients with neurofibromatosis type 1. J Neurosurg 118, 142–148 (2013).

12. Gutmann, D.H. et al. Neurofibromatosis type 1. Nat Rev Dis Primers 3, 17004 (2017).

13. Evans, D.G.R., Huson, S.M. & Birch, J.M. Malignant peripheral nerve sheath tumours in inherited disease. Clin Sarcoma Res 2, 17 (2012).

14. Lee, W. et al. PRC2 is recurrently inactivated through EED or SUZ12 loss in malignant peripheral nerve sheath tumors. Nat Genet 46, 1227–1232 (2014).

15. Pemov, A., Li, H., Presley, W., Wallace, M.R. & Miller, D.T. Genetics of human malignant peripheral nerve sheath tumors. Neurooncol Adv 2, i50–i61 (2020).

16. Lyskjaer, I. et al. H3K27me3 expression and methylation status in histological variants of malignant peripheral nerve sheath tumours. J Pathol 252, 151–164 (2020).

17. Nik-Zainal, S. et al. The life history of 21 breast cancers. Cell 149, 994–1007 (2012).

18. Bolli, N. et al. Heterogeneity of genomic evolution and mutational profiles in multiple myeloma. Nat Commun 5, 2997 (2014).

19. Kaufmann, T.L. et al. MEDICC2: whole-genome doubling aware copy-number phylogenies for cancer evolution. Genome Biol 23, 241 (2022).

20. Laks, E. et al. Clonal Decomposition and DNA Replication States Defined by Scaled Single-Cell Genome Sequencing. Cell 179, 1207–1221 (2019).

21. Fehrmann, R.S. et al. Gene expression analysis identifies global gene dosage sensitivity in cancer. Nat Genet 47, 115–125 (2015).

22. Aibar, S. et al. SCENIC: single-cell regulatory network inference and clustering. Nat Methods 14, 1083–1086 (2017).

23. Gavish, A. et al. Hallmarks of transcriptional intratumour heterogeneity across a thousand tumours. Nature 618, 598–606 (2023).

24. Cao, S. et al. Estimation of tumor cell total mRNA expression in 15 cancer types predicts disease progression. Nat Biotechnol 40, 1624–1633 (2022).

25. Ma, Y. & Zhou, X. Spatially informed cell-type deconvolution for spatial transcriptomics. Nat Biotechnol 40, 1349–1359 (2022).

26. Baker, T.G. et al. Near haploidization is a genomic hallmark which defines a molecular subgroup of giant cell glioblastoma. Neurooncol Adv 2, vdaa155 (2020).

27. Cortes-Ciriano, I. et al. Genomic Patterns of Malignant Peripheral Nerve Sheath Tumor (MPNST) Evolution Correlate with Clinical Outcome and Are Detectable in Cell-Free DNA. Cancer Discov 13, 654–671 (2023).

28. Noble, R. et al. Spatial structure governs the mode of tumour evolution. Nat Ecol Evol 6, 207–217 (2022).

29. Fu, X. et al. Spatial patterns of tumour growth impact clonal diversification in a computational model and the TRACERx Renal study. Nat Ecol Evol 6, 88–102 (2022).

30. Li, H. & Durbin, R. Fast and accurate short read alignment with Burrows-Wheeler transform. Bioinformatics 25, 1754–60 (2009).

31. McKenna, A. et al. The Genome Analysis Toolkit: a MapReduce framework for analyzing next-generation DNA sequencing data. Genome Res 20, 1297–303 (2010).

32. Zheng, G.X. et al. Massively parallel digital transcriptional profiling of single cells. Nat Commun 8, 14049 (2017).

33. Li, H. et al. The Sequence Alignment/Map format and SAMtools. Bioinformatics 25, 2078–9 (2009).

34. Kaminow, B., Yunusov, D. & Dobin, A. STARsolo: accurate, fast and versatile mapping/quantification of single-cell and single-nucleus RNA-seq data. Biorxiv, 2021.05. 05.442755 (2021).

35. Browning, B.L., Tian, X., Zhou, Y. & Browning, S.R. Fast two-stage phasing of large-scale sequence data. Am J Hum Genet 108, 1880–1890 (2021).

36. Nilsen, G. et al. Copynumber: Efficient algorithms for single- and multi-track copy number segmentation. BMC Genomics 13, 591 (2012).

37. Benjamin, D. et al. Calling somatic SNVs and indels with Mutect2. BioRxiv, 861054 (2019).

38. Gori, K. & Baez-Ortega, A. sigfit: flexible Bayesian inference of mutational signatures. Preprint at bioRxiv, 10.1101/372896 (2018).

39. Tate, J.G. et al. COSMIC: the Catalogue Of Somatic Mutations In Cancer. Nucleic Acids Res 47, D941–D947 (2019).

40. Van Loo, P. et al. Allele-specific copy number analysis of tumors. Proc Natl Acad Sci U S A 107, 16910–5 (2010).

41. Chen, T. & Guestrin, C. Xgboost: A scalable tree boosting system. in Proceedings of the 22nd acm sigkdd international conference on knowledge discovery and data mining 785-794 (2016).

42. Stuart, T. et al. Comprehensive Integration of Single-Cell Data. Cell 177, 1888–1902 e21 (2019).

43. Patel, A.P. et al. Single-cell RNA-seq highlights intratumoral heterogeneity in primary glioblastoma. Science 344, 1396–401 (2014).

44. Van de Sande, B. et al. A scalable SCENIC workflow for single-cell gene regulatory network analysis. Nat Protoc 15, 2247–2276 (2020).

